# When conflicts get heated, so does the planet: social-climate dynamics under inequality

**DOI:** 10.1101/2020.09.15.298760

**Authors:** Jyler Menard, Thomas M. Bury, Chris T. Bauch, Madhur Anand

**Affiliations:** Department of Physics and Astronomy, University of Waterloo, Waterloo, Ontario, Canada; Department of Applied Mathematics, University of Waterloo, Waterloo, Ontario, Canada; School of Environmental Sciences, University of Guelph, Guelph, Ontario, Canada

## Abstract

Climate dynamics are inextricably linked to processes in social systems that are highly unequal. This suggests a need for coupled social-climate models that capture pervasive real-world asymmetries in the population distribution of the consequences of anthropogenic climate change and climate (in)action. Here we develop a simple social-climate model with group structure to investigate how anthropogenic climate change and population heterogeneity co-evolve. We find that greater homophily and resource inequality cause an increase in the global peak temperature anomaly by as much as 0.7°C. Also, climate change can structure human populations by driving opinion polarization. Finally, climate mitigation achieved by reducing the cost of mitigation measures paid by individuals tends to be contingent upon socio-economic conditions, whereas policies that achieve communication between different strata of society show climate mitigation benefits across a broad socio-economic regime. We conclude that advancing climate change mitigation efforts can benefit from a social-climate systems perspective.

## Main

The Yellow Vest movement, protests during the Greek economic crisis, and players of the Ultimatum Game each reveal how asymmetrical costs and benefits fuel resistance to decision-making outcomes. In the Yellow Vest movement, protesters clearly stated their desire to mitigate climate change but would not support measures that unfairly impacted the working class [1, 2], while real-world players of the Ultimatum game will not accept offers deemed unfair, even when the Nash equilibrium predicts they should do so [3, 4, 5, 6].

Such asymmetries are also pervasive in climate change mitigation and the consequences of climate (in)action, such as: (1) asymmetry of contribution to the problem, wherein those who have benefited most from industrialization have contributed most to causing a climate emergency; (2) asymmetry of impact, wherein the worst consequences of climate change will fall on those least responsible; (3) asymmetry of power or representation, wherein the most affected do not always have the loudest or most heard voice; and (4) the asymmetry of responses to climate change, wherein some groups may be left behind during the transition to a low-carbon economy [7]. For instance, the Yellow Vest movement exemplifies how austerity measures applied to those least responsible and least powerful can exacerbate resource inequality and thereby lead to dissatisfaction.

Models increasingly help us understand the interactions between the carbon cycle, the climate system, and human processes, and the impact of policies [8, 9]. To ensure those policy decisions are robust to uncertainties, multiple scenarios are often laid out, ranging from carbon emission trajectories (RCPs) [8, 10] to socioeconomic systems pathways (SSPs) [11]. Influence in these models usually flows in one direction, from socioeconomic systems to the Earth system. Yet, two-way feedback mechanisms link climate and social processes: human behaviour changes the climate [12], and the climate changes human opinions [13, 14] and consequently human behaviour. Coupled human-environment models are already widely applied to other study systems such as fisheries and forests [15, 16, 17, 18, 19, 20, ?] and the need for coupled social-climate models has been noted [21, 22, 23]. Some integrated assessment models (IAMs) address this limitation by linking economic and climate systems [24, 25], but IAMs tend to omit non-market transactions, information asymmetries [24, 26], and feedbacks both within social systems and between social and Earth systems.

The need for simple social-climate models to better develop intuition and understanding of when social processes are important has been raised in the literature [9, 21]. Elsewhere, it has been suggested that evolutionary game theory may be an effective way of modelling social mechanisms as a dynamic process coupled to climate dynamics [23]. Recent coupled social-climate models have explored the implications of emergent social-climate dynamics [27, 21, 28, 22, 23], including the role of our reaction to extreme climate events where human behaviour is described by the theory of planned behaviour [27], and the role of social learning [28] and social norms [27, 28], in determining climate trajectories. We furthermore suggest that the asymmetries engendered by having distinct groups with different resource levels could play a significant role in social-climate dynamics and should be addressed in models.

Here we develop a coupled social-climate model to focus on the asymmetry of impact in the form of resource inequality and how austerity-induced dissatisfaction interacts with both climate mitigation efforts and climate system dynamics. Our objectives are (1) to investigate how population heterogeneity, homophily, and dissatisfaction affect the global average temperature anomaly predicted by an Earth system model; (2) to show how social-climate modelling may provide insights into climate change mitigation against a backdrop of powerful social forces; and (3) to introduce a method of modelling coupled social-climate systems with heterogeneous social structure.

## Model overview

To address our objectives we (1) use evolutionary game theory to develop a coupled social-climate model with heterogeneous population structure; (2) use the model to investigate how population heterogeneity changes social-climate dynamics; and (3) explore how the peak temperature anomaly responds to potential social-climate mitigation pathways across the parameter space. Our model is intended to gain intuition for nonlinear interactions between climate systems and heterogeneous social systems, rather than providing an empirically validated projections for policy-making. Hence, we opted to develop a minimal model. Details of our model appear in Methods.

The population consists of two groups differing with respect to (1) population size, (2) how costly it is for individuals to adopt a climate mitigation strategy (3) baseline resources available to individuals in the group, and (4) the extent to which the climate change impacts their resources. These differences are intended to capture relevant socio-economic inequalities within populations [7] and we refer to these features collectively as population heterogeneity. We considered a population divided into a ‘rich’ (R) and a ‘poor’ (P) group (Figure 1, left box). Social dynamics within each group are governed by imitation (social learning) dynamics, whereby individuals learn the opinions of their peers and change their opinions according to the group-specific utility functions. Individuals may adopt one of two opinions: mitigate climate change, or do not mitigate climate change. The utility functions governing decision-making depend on several factors, including social norms within the group, the perceived cost of climate change, the perceived cost of mitigation and, in the case of the poor group, a dissatisfaction term. Interaction between the two groups occurs via imitation dynamics weighted by homophily in both groups, and dissatisfaction amongst the poor group. The extent to which individuals imitate and make decisions based on social norms in the other group is governed by the homophily factor *h.* At the extreme values, *h* = 1 recovers no homophily (equal imitation within and across groups), and *h* = 0 resembles pure homophily (imitation only within each group). The dissatisfaction term captures the negative effect of resource inequality and inaction of the rich group on the poor group’s incentive to mitigate.

**Figure 1:**
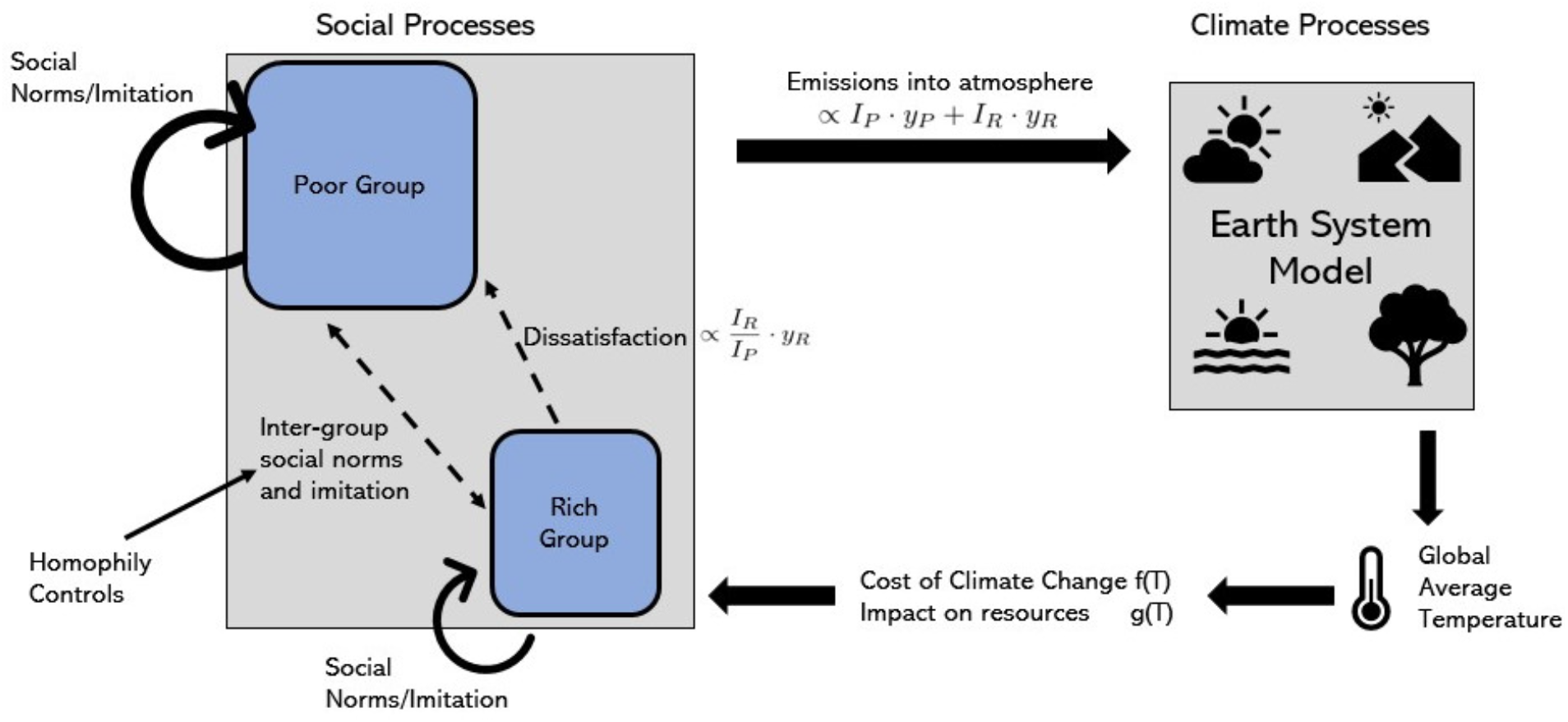
Schematic diagram of coupled social-climate model. The climate system (right box) impacts the social system (left box) via global average temperature (T). The social system impacts the climate system via carbon emissions from non-mitigators in the rich (R) and poor (P) groups. Emissions from group *g* are proportional to the fraction of non-mitigators (*y_g_*) and the group resource level (*I_g_*). Homophily controls the amount of imitation and influence of social norms between the groups. Social behaviour in the poor group is influenced by a dissatisfaction term, which is proportional to the level of inequality (ratio of rich to poor resource level) and fraction of non-mitigators in the rich group.

The climate model (right side of Figure 1) is a simple Earth System Model [29] consisting of a carbon cycle and a greenhouse effect. Emissions from the population enter the atmosphere, causing a greenhouse effect, and are then processed through the carbon cycle. Warming due to the greenhouse effect impacts the social dynamics through perceived dangers of climate change (an increase in projected future climate change based on recent trends increases the incentive to mitigate), and through impacting each group’s resources (to different extents – the rich group feels less impact compared to the poor group).

## Results

### Homophily increases the global peak temperature anomaly

We find that homophily increases the global peak temperature anomaly by slowing the spread of mitigative behaviour between groups via imitation (Eq. 2) and social norms (Eq. 1). We generated plots of the global temperature anomaly (*T*, blue), and proportion of poor (*x_P_*, red) and rich (*x_R_,* black) mitigators versus time, for various values of the strength of homophily (*h*) and the relative importance of dissatisfaction with resource inequality and mitigation efforts by the rich versus climate change impacts (*d/f_max_*) in the decision-making process (Figure 2). Across the full range of values for *d/f_max_,* stronger homophily always increases the peak temperature anomaly by approximately 0.3°*C*. When homophily is non-existent (*h* = 0), mitigative behaviour spreads readily between the rich and poor group on account of imitation and social norms, hence both groups adopt widespread mitigation on a rapid and roughly comparable timescale, although the rich group leads the adoption of mitigation when dissatisfaction is high, since the poor group is unwilling to pay the cost of mitigation unless the rich group also does so. On the other hand, when homophily is high (*h* = 1) it prevents the spread of mitigative behaviour between groups and therefore they adopt mitigation at very different timescales, depending on the strength of dissatisfaction. When dissatisfaction is strong (large *d/f_max_*), the poor group never adopts mitigation over the simulation time horizon. Whereas, when dissatisfaction is weak (small *d/f_max_*), it is the rich group that lags behind on mitigation since the poor group is stimulated to take action on climate mitigation on account of experiencing greater climate impacts. But in both cases, the lack of between-group imitation means that rich and poor groups adopt mitigation to very different degrees and thus the peak global temperature anomaly is higher.

**Figure 2:**
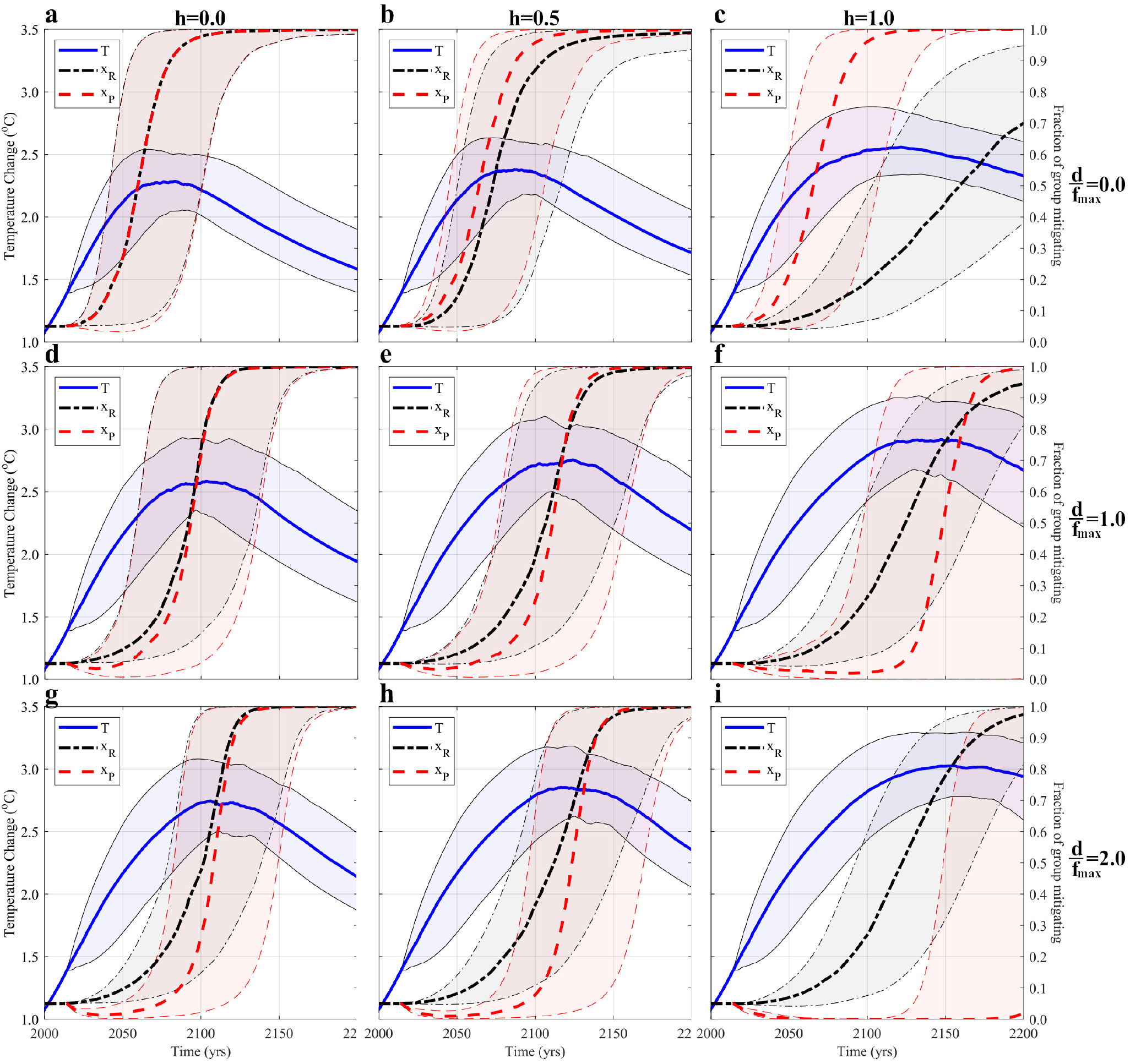
Homophily prevents the spread of mitigative behaviour and increases the peak temperature anomaly. Median trajectories with 95% confidence intervals (CI) of global temperature (blue solid) and proportion of mitigators for the rich (black dash-dotted) and poor (red dashed) groups. Panels show differing values of homophily (*h*) and dissatisfaction (*d*), with homophily increasing from left to right and dissatisfaction increasing from top to bottom. Median values are computed from 100 simulations, with parameter values (excluding those shown) drawn from triangular distributions defined in SI Text Table 2.

### Climate system feedback drives population polarization

The strong divergence of opinions between rich and poor groups when homophily is high (*h* = 1, Figure 2) represents a population polarization. However, there are two different mechanisms behind the polarization. The mechanism in operation depends upon the strength of dissatisfaction relative to climate change impacts (*d/f_max_*) in the decision-making process, even though the end result is always a higher temperature anomaly. When dissatisfaction is less important than climate impacts (*d/f_max_* < 1), the poor groups adopts mitigation faster, but homophily partly counteracts the unifying effect of social norms and slows the spread of mitigative behaviour from the poor group to the rich group (Figure 2abc). The mechanism at play here is simply different groups acting independently according to the socio-economic parameters governing their best interest, such that the poor group chooses mitigation rapidly while the rich group lacks behind.

In contrast, when *d/f_max_* > 1, the rich group adopts climate change mitigation but the poor group is much slower to follow-a “Yellow Vest” effect. Initially, the poor group is unwilling to mitigate until the rich group takes initiative. When the rich group finally adopts mitigation, dissatisfaction declines as a result (Eq. 5), but strong homophily slows the spread of mitigative behaviour from the rich group to the poor group (Figure 2ghi, SI Text Figure S9bc). This problem is compounded by the fact that the initial lack of mitigative behaviour in the rich group causes mitigation in the poor group to decline to very low levels in the early portion of the model trajectory, making it a strong social norm not to mitigate (Figure 2ghi). In the case where homophily is also strong (*h* = 1, Figure 2i), by the time mitigation becomes established in the rich group, there are so few mitigators in the poor group and so few opportunities for cross-group interaction, that mitigative behaviour recovers only very weakly by the end of the time horizon in most of the model realizations. As a result of dissatisfaction suppressing the timely adoption of mitigation by the poor group, the global temperature anomaly peaks at 3.0°*C*. The mechanism at play here is a strong synergy between the negative effects of homophily, norms and dissatisfaction.

The emergence of a polarized population under both mechanisms occurs because climate feedback structures the human population. Our model posits that anticipated climate change induces a population response, and if the population is heterogeneous with respect to socio-economic parameters governing their reaction to climate change, climate change will first trigger a reaction in the groups that are predisposed to respond to it on account of their socio-economic conditions. This reaction will generate both real and anticipated long-term climate change mitigation that in turn reduces the perceived urgency for the other groups to act, which were already predisposed not to act. Hence there is an inherent tendency for climate feedback to generate a polarized population. We suggest that this is a specific example of a more general mechanism for environmental systems to structure opinion and behaviour in human populations that have preexisting heterogeneity with respect to socio-economic privilege.

### Asymmetric impacts induce polarization despite lack of homophily

Surprisingly, symmetries in the impact of climate change on each group’s resources can increase peak polarization between groups even in the absence of homophily. When the rich group’s resources are not very affected by climate change, intensifying the impact on the poor group’s resources increases peak polarization (Figure 3c); additionally, increasing how abruptly the poor group’s resources are impacted does, as well (Figure 3d). Both of these results are driven by either increasing the nonlinearity or the maximum value of Eq. 6 thereby translating into increased dissatisfaction (Eq. 5), which directly translates into reduced incentive to mitigate in the poorer group (Eq. 1). Because the effect of dissatisfaction and resource inequality affect primarily the poorer group, increasing the size of the poorer group results in a significant increase in peak temperature anomaly (Figures S5, S6, and SI Text Figure S7), while shifting people towards the richer group has a much smaller effect on decreasing the peak temperature anomaly. The asymmetry in effect occurs because dissatisfaction can disincentivize a larger proportion of the total population from mitigating (SI Text Figure S7). Decreasing homophily can mitigate the impact of dissatisfaction (SI Text Figure S7), because then mitigative behaviour can spread between groups. This causes the rich group to adopt it faster, directly reducing the amount of dissatisfaction in the poor group (Eq. 5)

**Figure 3:**
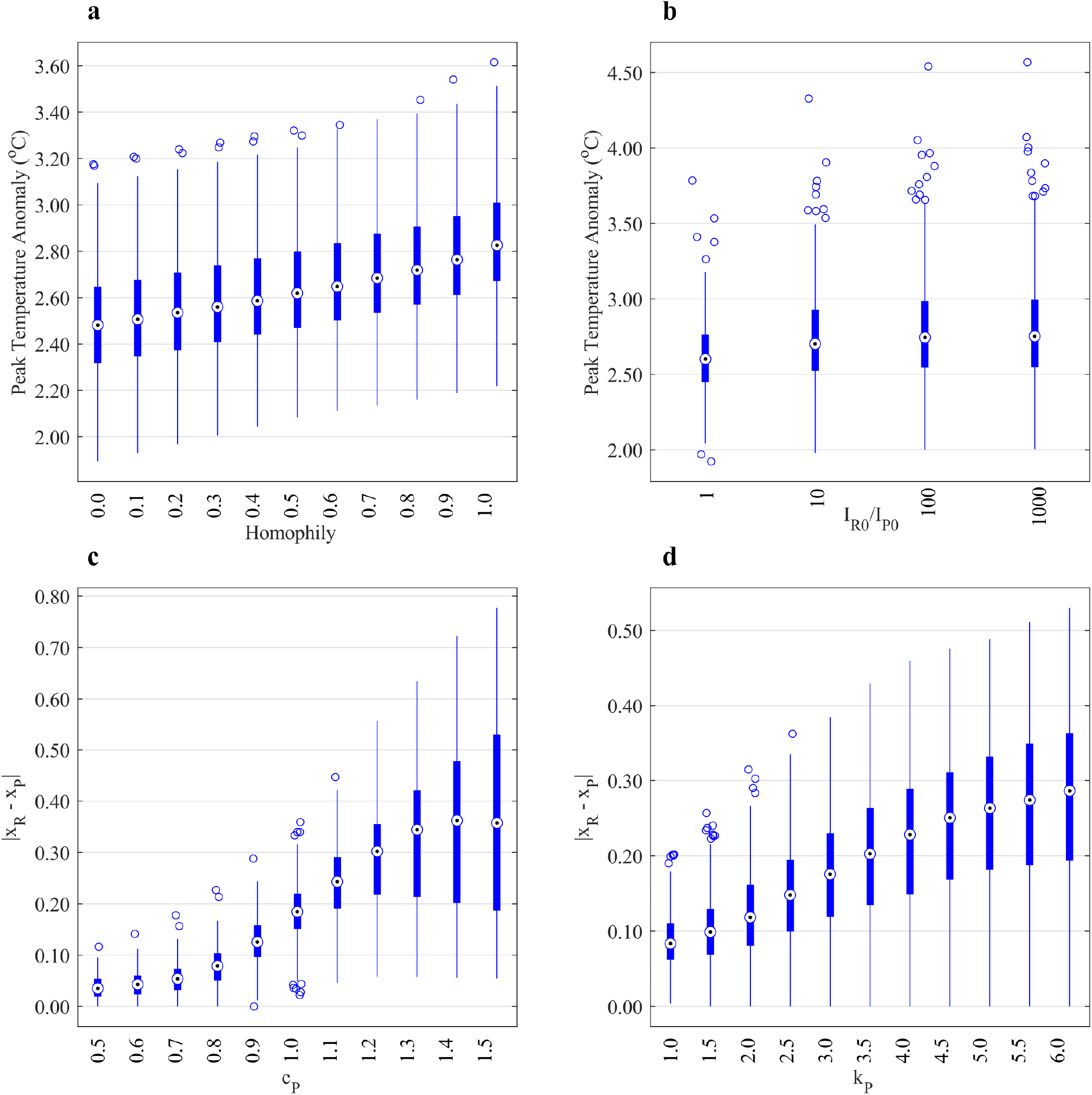
Intense asymmetries in climate change impact on resources can induce opinion polarization. Peak temperature anomaly versus (a) homophily and (b) relative initial resource level, and peak strategy difference versus (c) how intense the poor group’s resources are affected by increasing global temperature (*c_P_*), and (d) how abruptly the poor group’s resources are affected (*k_P_*). Boxplots report 500 simulations of the model with parameters sampled from triangular distributions as defined in Table 2 (SI Text). Plot (a) suggests increasing homophily increases peak temperature anomaly, while plot (b) shows very little relationship between initial resource inequality and peak temperature anomaly (with the exception of more extreme outliers). Plots (c) and (d) assume zero homophily, and suggest that climate-driven resource inequality can drive social polarization in our model.

However, that assumes the relative importance of dissatisfaction and climate change impacts are of similar magnitude; i.e. 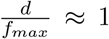. Yet, we find a monotonic relationship between 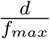 and peak polarization (Figures 4b, S5b). When the intensity of dissatisfaction is less, peak polarization is also less, leading to a reduced peak temperature anomaly (Figures 4a, S5a). Conversely, increasing the importance of dissatisfaction increases peak polarization, leading to an increased peak temperature anomaly. Interestingly, a larger peak polarization appears to have a saturating effect on the peak temperature anomaly. The dominant reason for this seems to be the rate at which the rich group becomes mitigative becomes independent of changes in 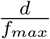 (SI Figure S10). Qnce dissatisfaction is much more salient to decisionmaking than climate outcomes 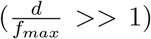 the poor group’s utility function is very negative – when the rich group is not mitigating – which pushes them towards non-mitigation. The rich group becoming mitigative reduces that dissatisfaction, allowing climate outcomes to become central to decision-making and the poorer group to become mitigative. This process is dependent on how quickly the rich group responds to climate change.

**Figure 4:**
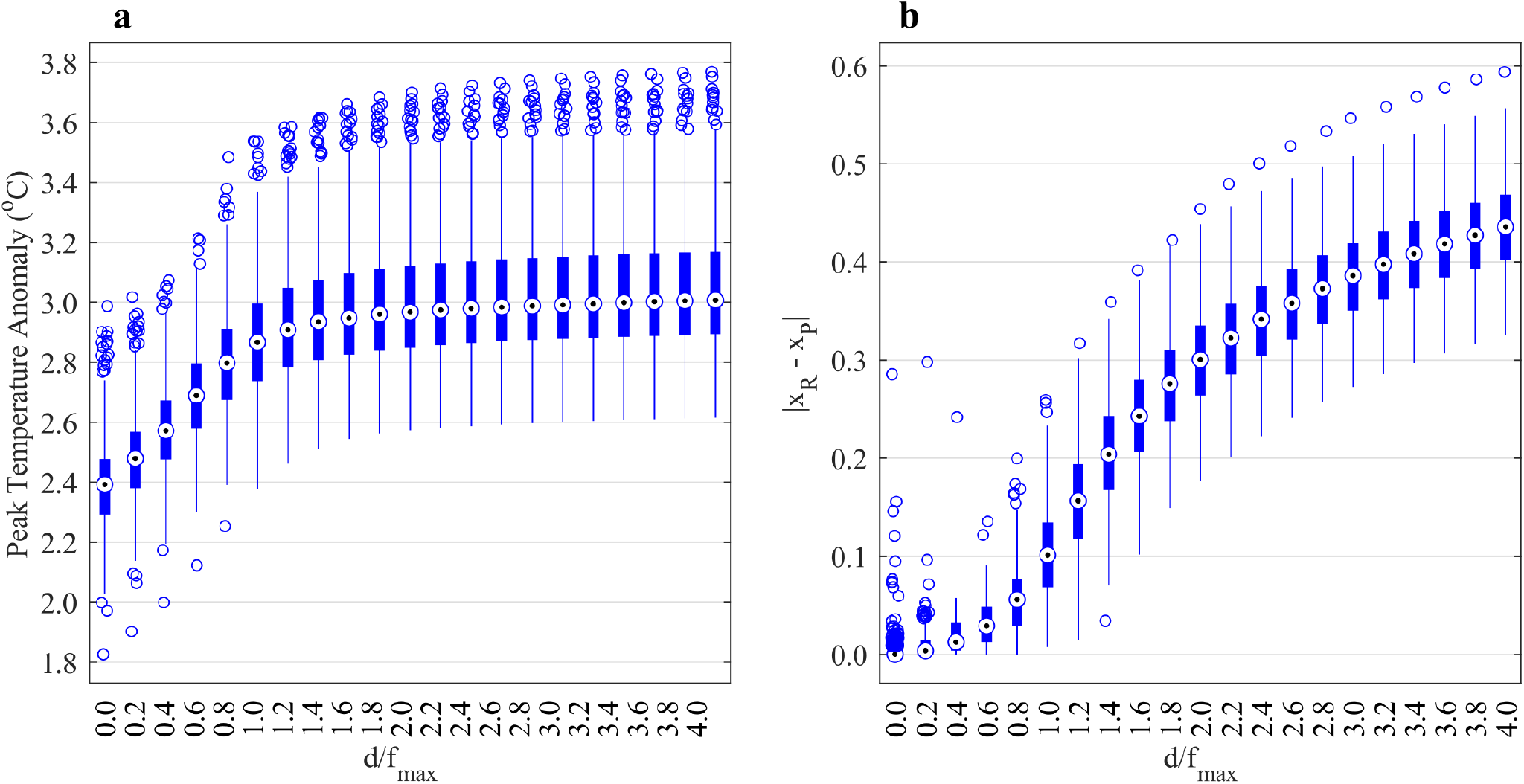
Increasing the relative importance of dissatisfaction versus climate change impacts increases opinion polarization and peak temperature anomaly. Box-and-whisker plots over 500 simulations for (a) peak temperature anomaly, and (b) peak polarization vs. ratio of dissatisfaction maximum to warming cost function maximum, at zero homophily and minimal impact on rich group’s resources. All other parameter values (excluding those shown) drawn from triangular distributions defined in SI Text Table 2.

### Sensitivity analysis

Univariate sensitivity analysis with tornado plots suggests climate outcomes depend considerably on social factors (*κ* and *ρ_R_*), the ability for the mitigative strategy to spread between groups (*h*), and the relative importance of dissatisfaction and the cost of climate change (*d, f_max_, T_c_*, and *d_c_*) (Figure S5). Varying dissatisfaction with resource inequality and inactivity in the rich group is for the poor group lowers the peak temperature anomaly by up to 0.5°*C* from the baseline model, and increases mitigative opinion to ≈80% of the total population, relative to when dissatisfaction is absent.

Interestingly, in the upper and lower limits of homophily, the parameters with highest sensitivity change (Figure S6). This is because in the absence of homophily the two groups respond to each other and the climate dynamics, while in full homophily they ignore each other and respond only to the climate. The strength of social norms (*δ*) becomes more impactful in the absence of homophily, tightening how strongly the opinion of one group is tied to norms of the other group. When homophily is at the other extreme, the strength of social norms becomes much less impactful because it only affects the groups independently of each other. Additionally, when homophily is full, uncertainties in climate parameters become more impactful on climate outcomes (*S*, *χ*, *A,* and *F*_0_ all increase); the two groups stop responding to each other and respond only to the changing climate, making its dynamics more impactful to when the population begins mitigating.

How quickly the cost increases (*d_c_*) influences the peak temperature anomaly by changing where the cost function is most sensitive to temperature. When it is lower, the cost of dissatisfaction quickly increases before the perceived cost of climate change can encourage mitigation, hence an increase in *T_c_* and a decrease in *d_c_* both increase the peak temperature anomaly (Figure S5, SI Figure S8)–the former because climate change is seen to be costly less quickly, and the latter because the poor group becomes non-mitigative more quickly.

### Socially-driven approaches to climate mitigation

Because social-climate models incorporate both social and climate processes, they provide a method to compare the impacts of social and economic policy interventions through the use of parameter planes, which allows plotting pathways to climate mitigation through pairwise parameter changes. This analysis suggests that climate change might be mitigated most effectively not by economic policies to make mitigative technologies more accessible (although this is still important), but rather by social policies designed to encourage communication between different strata of society.

This prediction is apparent in parameter planes governing the strength of social norms (*δ*), cost of mitigation (*α*_*R*0_ and *α*_*P*0_), amount of homophily (*h*), and the importance of dissatisfaction relative to climate impacts (*d/f_max_*) (Figure 5). For instance, simultaneously increasing social norms and decreasing homophily strongly reduces the peak temperature anomaly (Figure 5d). This is because a stronger social norm maintains group cohesion, so when homophily is also low, both groups converge to the same opinion. In contrast, homophily has little impact when social norms are weak because social norms are one of the main routes on which homophily operates in our model.

**Figure 5:**
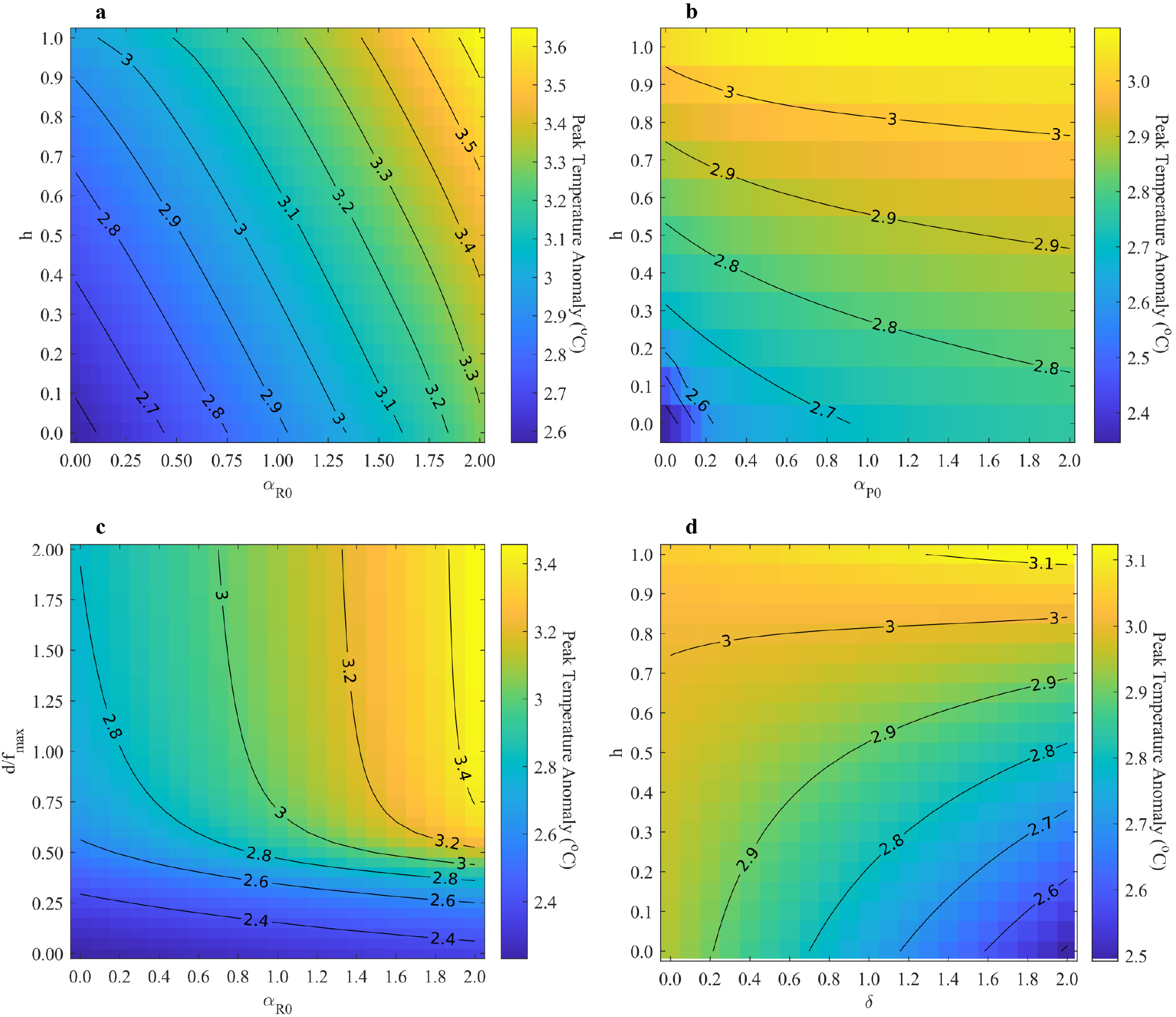
Social pathways to climate mitigation. Contour plots showing peak temperature anomaly attained at specific values of: (a), homophily and cost of mitigation for the rich group (*α*_*R*0_); (b), homophily and cost of mitigation for the poor group (*α*_*P*0_); (c), relative cost of dissatisfaction to perceived cost of climate change 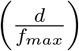, and *α*_*R*0_; and (d), homophily and strength of social norms (*δ*). All other parameters are held at baseline values (see SI Text for additional contour plots). Reducing the amount of homophily reduces peak temperature anomaly, and increases the effectiveness of drecreases costs of mitigation.

Similarly, reducing homophily, reducing dissatisfaction, or increasing the anticipate impacts of climate change show significant benefits across a broad region of parameter space, whereas the benefits of reducing the cost of mitigation are either weaker, or more contingent on the social parameter values (Figure 5abd). For instance, reducing the cost of mitigation for the rich group tends to decrease the peak temperature anomaly regardless of the amount of homophily, but not if dissatisfaction is low in the poor group and/or if anticipated climate impacts are high (Figure 5ac). This is because the primary mechanism by which reducing *α*_*R*0_ benefits climate change is through uptake of mitigation by the rich group reducing dissatisfaction among the poor group and thus enabling the (more numerous) poor group to adopt mitigation. When dissatisfaction is small (or conversely, anticipated climate change impacts are very large), this mechanism is not necessary to mitigate climate change. Unexpectedly, our model predicts that reducing reducing *α*_*P*0_ does very little to reduce the peak temperature anomaly, unless we willing to make mitigation nearly costless for the poor group, and to reduce homophily to extremely small values (Figure 5b).

## Discussion

We introduced a social-climate model based on asymmetric evolutionary game theory with a greater amount of population heterogeneity than existing models. Among other findings, our model predicted that climate feedbacks can structure human populations by driving opinion polarization. Our model also predicted that the benefits of economic policies making mitigation less costly are often contingent upon social and economic conditions, whereas social policies designed to encourage communication between different strata of society exhibit more robust benefits across a broad range of socio-economic conditions.

Political polarization and politics surrounding group identity are growing issues. Digital social media enables individuals to self-sort into homophilous echo chambers. And, the “Yellow Vest” movement suggests dissatisfaction with the perceived unwillingness of more resourceful groups to mitigate emissions is one of many factors that can also lead to polarization. Our model demonstrates how homophily, in conjunction with climate-exacerbated dissatisfaction, can lead to polarization in climate responses. This is just one mechanism by which a worsening climate can deepen or magnify existing problems within social systems [7]. Conversely, because human behaviour then impacts the climate, corroded social systems have the potential to worsen climate anomalies, as our model illustrates. Increased population heterogeneity, homophily, and dissatisfaction can all lead to an increased global temperature anomaly. A worrisome feedback loop thereby suggests itself: more extreme temperature anomalies worsen social systems thereby worsening temperature anomalies. As a human-environment system, social-climate dynamics may warrant more investigation, and may provide an additional method of incorporating work done by social scientists and humanities scholars [30].

The factors by which dissatisfaction becomes dominant are fourfold: (1) homophily disrupts intergroup learning, (2) climate-driven resource inequality creates resentment, (3) lack of action by the resource-rich group generates dissatisfaction, and (4) a small perceived risk of climate change makes the dissatisfaction relatively more important for decision-making. There may be solutions to address each of these mechanisms. For (1), there is some evidence intergroup contact can reduce homophily and lead to spreading of environmental concern [31]; (2) and (3) may be addressed through carbon tax redistribution programs [32], or investments in entrepreneurship and certification programs (to facilitate career change); and (4) may be addressed through increased speed and reliability of attributing extreme events – and other local changes – to climate change.

We opted for a minimal model that nonetheless incorporated a selection of realistic social features such as population heterogeneity and social learning. However, the model’s simplifying assumptions could influence its predictions. Many of our parameters are not directly measurable in experiments or surveys (although they could be inferred by curve-fitting). Our resource model ignores time dependencies. We assume only two groups and only two strategies with respect to climate action. These limitations diminish the usefulness of our model for accurate social-climate projections. However, as noted in [27, 21, 28, 22, 23], simple models are still valuable tools to understand processes, attempt new modelling methods, and to explore how the approach could be effective in suggesting policy changes [9]. To this end, our results suggest that social-climate interactions in a heterogeneous population can have nontrivial impacts on both climate trends and opinion dynamics, and thus merit further study. As climate change is driven by social and economic processes we should not forget that climate change is ‘everything change’ – and will require a complementary social change in order for us to mitigate it [33, 34].

## Methods

According to our objectives we used a relatively simple coupled social-climate model. The benefits of this approach are threefold: (1) it is less computationally demanding; (2) its behaviour is easier to understand; (3) it is more tractable to analyse. A conceptual diagram of the model appears in Figure 1 and we explain each of the components in the following subsections.

### Software

We used Matlab R2020a to implement our delay-differential equation model, and used fourth-order Runge-Kutta to solve the initial boundary value problem.

### Social Model

#### Social learning and homophily

We model social dynamics using asymmetric evolutionary game theory [35, 36]. There are two ways of making a game asymmetric: give players different strategy sets, or give players different utilities for using the same strategies. We focus on the latter. Individuals can follow either a mitigation (M) or a non-mitigation (N) strategy [28]. The M strategy corresponds to mitigating carbon emissions released to the atmosphere, while the N strategy does not reduce emissions. Unlike Ref. [27, 28], individuals are in one of two groups, which we label as ‘rich’ (R) and ‘poor’ (P). Since the two groups can use the same strategies but get different utility, we model an asymmetric evolutionary game.

Specifically, the utility of being a mitigator or non-mitigator in each group is determined by several factors: social norms within each group, the perceived cost of climate change, the perceived cost of mitigation, and in the case of the poor group, a dissatisfaction term. The utility of being a poor mitigator is

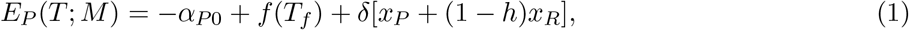

where *f*(*T_f_*) is the perceived cost of the temperature anomaly *T_f_* [28]; *δ* is the strength of social norms [37, 38]; *x_P_* (*x_R_*) is the proportion of mitigators in the poor (rich) group, and *h* is the homophily parameter (described in more detail below). The remaining utility functions are in the supplementary information. Differences between the utility function above (Eq. 1) and the utility functions reported in the supplementary info are: for the richer group there is no cost of dissatisfaction term; and, the utility of being a non-mitigator does not include a ‘cost of mitigating/non-mitigating’ because they could be absorbed into the net cost parameter. Normalizing the social norms term in the utility function is shown in the supplementary information to have no affect on the main results.

Social learning is implemented by modifying imitation dynamics described in previous publications [28, 35, 36], but individuals can have differing receptivity to information from individuals belonging to different groups. Hence, following [39] we introduced a homophily parameter *h* ∈[0,1] controlling the extent to which the rich and poor groups imitate, and exert social pressure on, each other. The imitation dynamics for the poor group is then

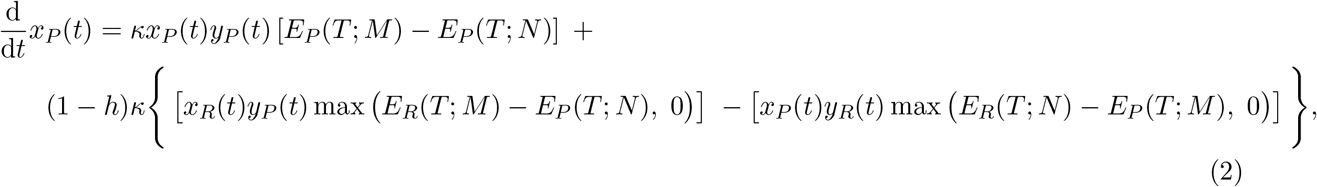

where *x*_*R*(*P*)_ is the proportion of rich (poor) mitigators, and *y*_*R*(*P*)_ is the proportion of rich (poor) non-mitigators. The dynamics for the rich group are equivalent with subscript “R” substituted for “P”. The derivation of equation 2 is explained more fully in SI Text. Heuristically, we can understand it as follows. The first term is the replicator equation, as described in [36, 35]; it describes imitation of opinions within a group. The second term describes imitation of strategies between groups and is scaled by the homophily parameter *h*. Hence, if individuals listen only to members of their own group (*h* = 1), they effectively see one population and we retrieve the social dynamics of [28].

#### Temperature Projection

Given that climate forecasts and individual experience influence perception of climate change [40], we assume that individuals extrapolate perceived future climate change linearly from recent trends using

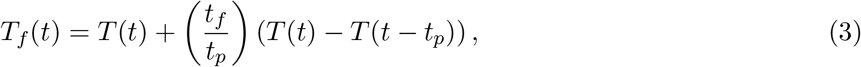

where *t_f_* is the number of years into the future to project, and *t_p_* is the number of years into the past.

#### Cost of Climate Change

To approximate nonlinear changes in the perceived cost of climate change we use a sigmoid function

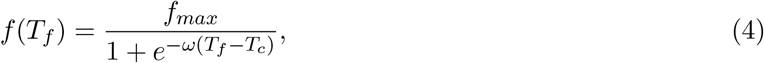

where *ω* varies the nonlinearity of the function, *f_max_* is the maximum cost of climate change, and *T_c_* is the temperature where the cost of climate change is most sensitive to change.

#### Dissatisfaction

We introduce a dissatisfaction term in the net cost of being a mitigator in the poor group

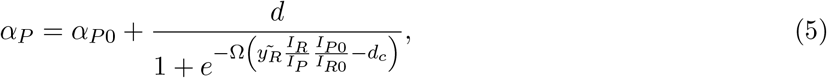

where 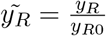 is the number of non-mitigators in the rich group normalized by the initial number of non-mitigators in the rich group. This is meant to capture dissatisfaction of the poor group with the inaction of rich group, analogous to the dissatisfaction with unfairness seen in the Ultimatum game [4, 5, 6]. Dissatisfaction increases when there are more non-mitigators in the rich group 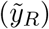, and when the the wealth gap between rich and poor is greater 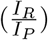. We used a multiplicative dissatisfaction 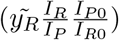 to ensure dissatisfaction is extensive: when the size of the rich group doubles at a given inequality, we expect the dissatisfaction to double. Increasing dissatisfaction increases the cost of mitigation, diminishing the incentive to mitigate climate change. For simplicity, we assume the cost of mitigation for the rich group is fixed.

#### Resource Inequality

The rich and poor groups each have an initial fixed resource endowment *I_R_*, *I_P_* with *I_P_* < *I_R_*. This resource endowment can be interpreted as relative income or relative influence in the sense that if a group has a large amount of resources it is able to have a larger impact. Each of the group’s resource endowment is affected by temperature change from the climate model according to

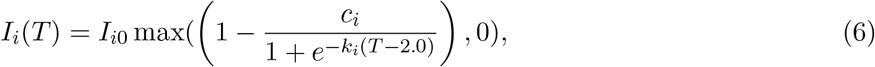

where *T* is the difference in temperature from initial. We define 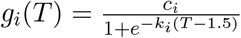 to be the ‘resource cost function’ and *i* ∈ {*R, P*}. *c_i_* is the maximum impact possible, and controls how extremely climate change impacts the group’s resources. *k_i_* controls the nonlinearity of, or how abruptly resources are impacted by, the response to changing temperature. In the baseline model, parameter intervals are chosen such that the resources of the rich group is always greater than that of the poor group, and the inequality between the two groups increases with increasing temperature anomaly to reflect how poorer countries and people will be worse affected relative to richer countries and people [7]. Resource level response to temperature anomaly is shown in the supplementary information (SI Figure S1)

In summary, imitating strategies between groups is affected by two new processes. The first is homophily, increasing its value decreases whether members of one group or wealth class listens to members of the other group. The second process is dissatisfaction. It controls whether members of the poor group deem whether the climate burden they bear is fair relative to the rich group’s willingness to also bear the climate burden. Dissatisfaction is influenced by the proportion of the rich group that is willing to participate in mitigation, and the inequality between the two groups. Resources are a third process and they contribute to whether dissatisfaction increases or not, and the extent to which a group has control over the carbon emissions.

#### Parameterization

Since our objective was qualitative analysis of model dynamics, we chose model parameters to obtain baseline dynamics that satisfied certain qualitative requirements. Many of the model parameters were chosen to ensure we are modelling the IPCC’s asymmetries of impact and power. The initial resource levels of each of the groups (*I*_*R*0_ > *I*_*P*0_), and the parameters controlling the nonlinearity and maximum of the income cost functions (*k_R_, k_P_*, *c_R_, c_P_*) were selected so that the rich group would have more resources than the poor group and that the poor group is worse affected by a changing climate. In the baseline model, the maximum of the dissatisfaction function was chosen so that it was the same as the maximum of the cost of climate change function. The reason for this is because we do not know exactly whether dissatisfaction with resource inequality, and with the inaction of a more resourceful group, is more important to decision-making compared to perceptions of climate change impacts.

### Climate Model

We use a simplified Earth System Model that captures each of the main aspects of the carbon cycle and a globally averaged greenhouse effect [29]. There are three primary reasons we selected this model: (1) it follows projections done by the current CMIP5 model when forced with the IPCC emissions scenarios (SI Text Figure 9); (2) it includes the carbon cycle and its effects on global warming, this is necessary because of how we are modelling social dynamics (mitigating or not mitigating atmospheric carbon emissions); (3) it is simple to use and implement, plus it has low computational cost. The model is from a previous publication [29] hence we describe it in more detail in the supplementary text. Here we describe the coupling between the Earth Systems Model and the social dynamics.

### Coupling of Social and Climate Model

In order to analyze the feedback effects of social processes on climate processes, we need to couple the climate model and social model. To do this, we scale carbon emissions with the proportion of non-mitigators (*y_i_*) and their resource level (*I_i_*). This amounts to saying that a larger proportion of non-mitigators will increase emissions. Because we are focused on asymmetries between two groups, we separate this forcing into two terms; one for the effect of the rich group and one for the poor group. This can be written as:

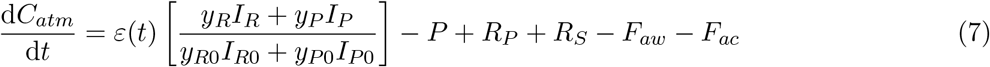

The first (second) term scaling *ε*(*t*) is the amount of atmospheric carbon emissions due to the non-mitigators in the rich (poor) group, normalized by total non-mitigator impact. We chose this normalization to ensure the relative impact of both the size of a group and their resource amount is included in their effect on emissions; e.g. if we had normalized each factor individually 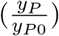, then we would erase the effect of the size of the group on emissions. The remainder of the climate model is included in SI Text.

## Acknowledgements

This research was supported by Natural Sciences and Engineering Research Council of Canada (NSERC) Discovery Grants and a New Frontiers in Research Fund (NFRF) Exploration Grant to MA and CTB.

## Author contributions

MA and CTB conceived of the study. All authors designed the research. JM analyzed the model and wrote the first manuscript draft. All authors edited the manuscript.

