## Supplementary Appendix for "When conflicts get heated, so does the planet: social-climate dynamics under inequality"

### 1.1 State variables

| Variable/Process | Definition | Unit |
|---|---|---|
| $x_R$ | proportion of mitigators in the rich group | 1 |
| $x_P$ | proportion of mitigators in the poor group | 1 |
| $C_{at}$ | deviation of atmospheric $CO_2$ (from pre-industrial 1800) | GtC |
| $C_{oc}$ | deviation of $CO_2$ in ocean | GtC |
| $C_{veg}$ | deviation of $CO_2$ in vegetation | GtC |
| $C_{so}$ | deviation of $CO_2$ in soil | GtC |
| $T$ | deviation in temperature | $^oC$ |
| $\varepsilon(t)$ | $CO_2$ emissions in absence of mitigation | GtC/yr |
| $P$ | carbon uptake from photosynthesis | GtC/yr |
| $R_{veg}$ | respiration from vegetation | GtC/yr |
| $R_{so}$ | respiration from soil | GtC/yr |
| $F_{oc}$ | flux of $CO_2$ from atmosphere to ocean | GtC/yr |

Table 1: **State variables and dynamic processes in the social-climate model**.

### 1.2 Parameter definitions

| Parameter | Definition | Baseline Values / Intervals | Unit | Source |
|---|---|---|---|---|
| $I_{R0}$ | initial income of rich group | (4.5, 5, 5.5) | 1 | - |
| $I_{P0}$ | initial income of poor group | (3.15, 3.5, 3.85) | 1 | - |
| h | homophily parameter | (0, 0.5, 1.0) | 1 | [39] |
| $\rho_R$ | size of of rich group | (0.10, 0.25, 0.40) | 1 | - |
| $c_R$ | maximum of income cost function $g_R(T)$ | (0.36, 0.4, 0.44) | 1 | - |
| $c_P$ | maximum of income cost function $g_P(T)$ | (0.765, 0.85, 0.935) | 1 | - |
| $k_R$ | nonlinearity of income impact function $g_R(T)$ | (0.9, 1.0, 1.1) | 1 | - |
| $k_P$ | nonlinearity of income impact function $g_P(T)$ | (1.35, 1.5, 1.65) | 1 | - |
| $\alpha_{P0}$ | cost of mitigation for poor group | (0.9, 1.0, 1.1) | 1 | - |
| $\alpha_{R0}$ | cost of mitigation for rich group | (0.45, 0.5, 0.55) | 1 | - |
| $d$ | maximum cost of dissatisfaction | (0.0, 5.0, 10.0) | 1 | - |
| $\Omega$ | nonlinearity of dissatisfaction cost | (2.7, 3.0, 3.3) | 1 | - |
| $d_c$ | critical value of dissatisfaction cost | (1.35, 1.5, 1.65) | 1 | - |
| $C_{at0}$ | initial $CO_2$ in atmosphere | (590, 596, 602) | GtC | [41, 29] |
| $C_{ao0}$ | initial $CO_2$ in ocean reservoir | $(1.4, 1.5, 1.6)\times10^5$ | GtC | [42] |
| $C_{veg0}$ | initial $CO_2$ in vegetation reservoir | (540, 550, 560) | GtC | [41, 29] |
| $C_{so0}$ | initial $CO_2$ in soil reservoir | (1480, 1500, 1520) | GtC | [41, 29] |
| $T_0$ | initial average atmospheric temperature | (288, 288.15, 288.3) | K | [29] |
| $k_P$ | photosynthesis rate constant | (0.175, 0.184, 0.193) | $\mathrm{yr}^{-1}$ | [41, 29] |
| $k_{MM}$ | photosynthesis normalising constant | 1.478 | 1 | [29] |
| $k_c$ | photosynthesis compensation point | $(26, 29, 32)\times10^{-6}$ | 1 | [43, 29] |
| $K_M$ | half-saturation point for photosynthesis | $(108, 120, 132)\times10^{-6}$ | 1 | [29] |
| $k_a$ | mole volume of atmosphere | $1.773 \times 10^{20}$ | moles | [44, 29] |
| $k_r$ | plant respiration constant | (0.0828, 0.092, 0.1012) | $\mathrm{yr}^{-1}$ | [41, 29] |
| $k_A$ | plant respiration normalising constant | $8.7039 \times 10^9$ | 1 | [29] |
| $E_a$ | plant respiration activation energy | (54.63, 54.83, 55.03) | kJ $\mathrm{mol}^{-1}$ | [45, 29] |
| $k_{sr}$ | soil respiration rate constant | (0.0303, 0.034, 0.037) | $\mathrm{yr}^{-1}$ | [41, 29] |
| $k_B$ | soil respiration normalising constant | 157.072 | 1 | [29] |
| $k_t$ | turnover rate constant | (0.0828, 0.092, 0.1012) | $\mathrm{yr}^{-1}$ | [41, 29] |
| $c$ | specific heat capacity of Earth's surface | $(4.22, 4.69, 5.16)\times10^{23}$ | $\mathrm{JK}^{-1}$ | [41, 29] |
| $a_E$ | Earth's surface area | $5.101 \times 10^{14}$ | $\mathrm{m}^2$ | universal |
| $\sigma$ | Stefan-Boltzmann constant | $5.67 \times 10^{-8}$ | $\mathrm{Wm}^{-2}\mathrm{K}^{-4}$ | universal |
| $L$ | latent heat per mole of water | 43 655 | $\mathrm{mol}^{-1}$ | universal |
| $R$ | molar gas constant | 8.314 | J $\mathrm{mol}^{-1}\mathrm{K}^{-1}$ | universal |
| $H$ | relative humidity | 0.5915 | 1 | calibrated |
| $A$ | surface albedo | (0.203, 0.225, 0.248) | $\mathrm{yr}^{-1}$ | [46, 29] |
| $S$ | solar flux | (1231, 1368 1504) | $\mathrm{Wm}^{-2}$ | [46, 29] |
| $\tau(CH_4)$ | methane opacity | (0.0208, 0.0231, 0.0254) | 1 | [47, 29] |
| $P_0$ | water vapor saturation constant | $(1.26, 1.4, 1.54)\times10^{11}$ | Pa | [29, 48] |
| $F_0$ | ocean flux rate constant | $(2.25, 2.5, 2.75)\times10^{-2}$ | $\mathrm{yr}^{-1}$ | [42] |
| $\chi$ | characteristic $CO_2$ solubility | (0.2, 0.3, 0.4) | 1 | calibrated |
| $\zeta$ | evasion factor | (40, 50, 60) | 1 | calibrated |
| $\kappa$ | social learning rate | (0.02, 0.05, 0.2) | $\mathrm{yr}^{-1}$ | [28] |
| $\delta$ | strength of social norms | (0.5, 1.0, 1.5) | 1 | [28] |
| $f_{max}$ | maximum of warming cost function $f(T)$ | (4, 5, 6) | 1 | [28] |
| $\omega$ | nonlinearity of warming cost function $f(T)$ | (1, 3, 5) | $\mathrm{K}^{-1}$ | [28] |
| $T_c$ | critical temperature of $f(T)$ | (2.4, 2.5, 2.6) | K | [28] |
| $t_p$ | # previous years used for temperature projection | 10 | yr | [28] |
| $t_f$ | # years ahead for temperature projection | (13.5, 15, 16.5) | yr | [28] |
| $s$ | half-saturation time for $\epsilon(t)$ from 2014 | (30, 50, 70) | yr | [28] |
| $\epsilon_{max}$ | maximum change in $\epsilon(t)$ from 2014 | (4.2, 7, 9.8) | GtC $\mathrm{yr}^{-1}$ | [28] |
| $x_0$ | initial proportion of mitigators | 0.05 | 1 | [28] |

Table 2: **Definitions and values for the parameters in the social-climate model.** Parameter values given as a tuple provide the lower bound, baseline, and upper bound values that are then used to define triangular distributions.

### 1.3 Resource level impacts

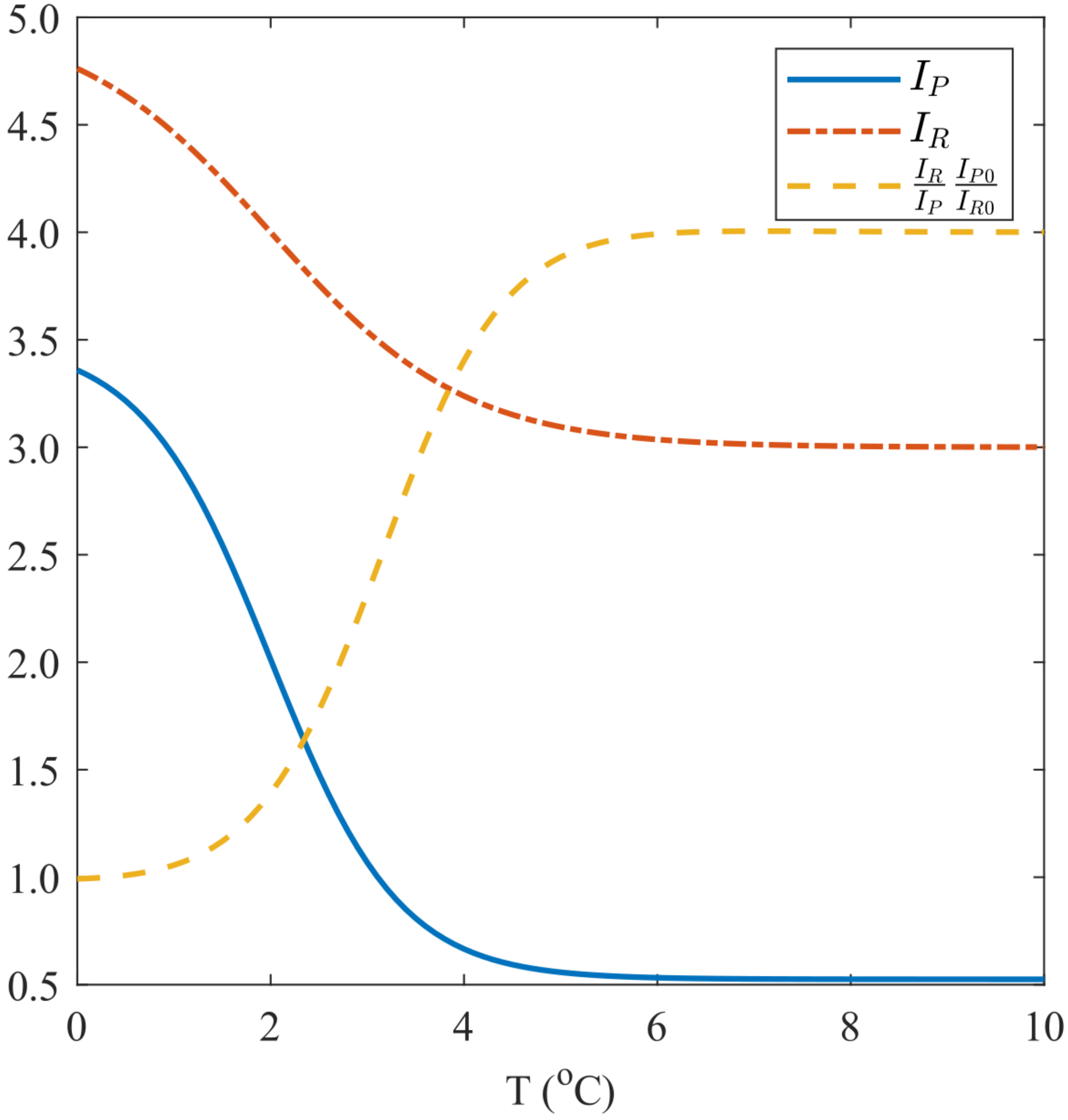


Figure S1: **Resource level with sigmoidal temperature impact function**. Baseline values are used for all parameters and are available in SI Text Table 2. Dashed line is the ratio $\frac{I_R}{I_P}\frac{I_{P0}}{I_{R0}}$. Dash-dotted (solid) line is the resource level of the rich (poor) group.

#### 1.4 Utility Functions

There are four payoffs:

$$\begin{aligned} E_P(T_f; M) &= -\alpha_P + f(T_f) + \delta P(M) \\ E_P(T_f; N) &= -f(T_f) + \delta P(N) \\ E_R(T_f; M) &= -\alpha_R + f(T_f) + \delta P(M) \\ E_R(T_f; N) &= -f(T_f) + \delta P(N) \end{aligned}$$

Where $\alpha_i$ is the net cost of mitigative behaviour for group $i$; $\delta$ is the strength of social norms, which is multiplied by the proportion of the population using the given strategy to reinforce said strategy; and, $\tilde{f}(T)$ is the cost of a temperature anomaly of T degrees. Including the effects of homophily, the four payoffs become:

$$\begin{aligned} E_P(T_f; M) &= -\alpha_P + f(T_f) + \delta[x_P + (1-h)x_R] \\ E_P(T_f; N) &= -f(T_f) + \delta[y_P + (1-h)y_R] \\ E_R(T_f; M) &= -\alpha_R + f(T_f) + \delta[(1-h)x_P + x_R] \\ E_R(T_f; N) &= -f(T_f) + \delta[(1-h)y_P + y_R] \end{aligned} \tag{8}$$

If homophily is not present ($h = 0$), then the proportions become the proportions of the total population using a given strategy. $\alpha_P$ depends on a dissatisfaction term, and is dependent on resource level of each of the groups. This is because lack of mitigative effort by other groups foments resentment that disinclines mitigative behaviour in the poor group. We make it easier for the richer group to contribute to mitigating climate change relative to poorer groups by having $\alpha_R < \alpha_P$ and also their resource level does not decrease as quickly or as significantly as in the poor group's.

#### 1.5 Normalized Social Norms

In the main text the social norms acts such that individuals in one group feel more pressure when $h = 0$ than when $h = 1$ because they perceive a larger population. This can be seen when we consider the payoffs where we substitute the maximum possible values of $x_R$, $y_R \in [0, \rho]$, $x_P$, $y_P \in [0, 1-\rho]$, substituting the maximums yields:

$$\begin{aligned} E_P(T_f; M) &\propto \delta[1 - \rho + (1-h)\rho] \\ E_P(T_f; N) &\propto \delta[1 - \rho + (1-h)\rho] \\ E_R(T_f; M) &\propto \delta[(1-h)(1-\rho) + \rho] \\ E_R(T_f; N) &\propto \delta[(1-h)(1-\rho) + \rho] \end{aligned}$$

so at maximum:

$$E_P(T_f; M) \propto \delta[1 - h\rho]$$
$$E_P(T_f; N) \propto \delta[1 - h\rho]$$
$$E_R(T_f; M) \propto \delta[1 - h + h\rho]$$
$$E_R(T_f; N) \propto \delta[1 - h + h\rho]$$

which shows that the maximum of the social norms term depends on $h$ and $\rho$. If we demand that it does not depend on $h$ or on $\rho$, we can divide by $[1 - h\rho]$ for the poor group and by $[1 - h + h\rho]$ for the rich group. Doing this social norm normalization does not change the main results, in fact in some cases it slightly furthers the trends seen in the main results, as follows.

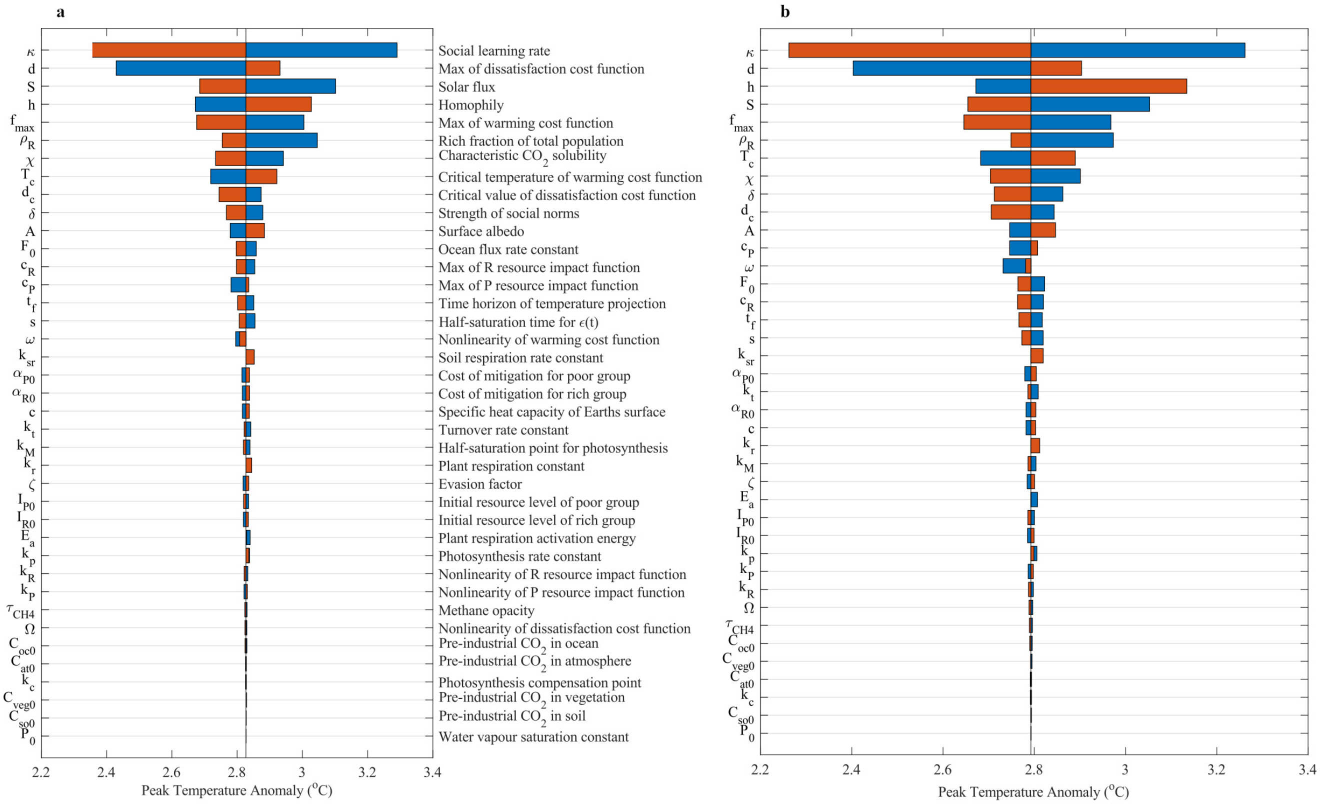


Figure S2: Comparison of (a) no normalization (main text), and (b) social norms normalized.

There are very few differences between the main text figure (left) and the tornado plot using the normalized social norms term (right). Homophily has a slightly higher relative importance, and the climate-related parameters drop in relative importance (but not by much).

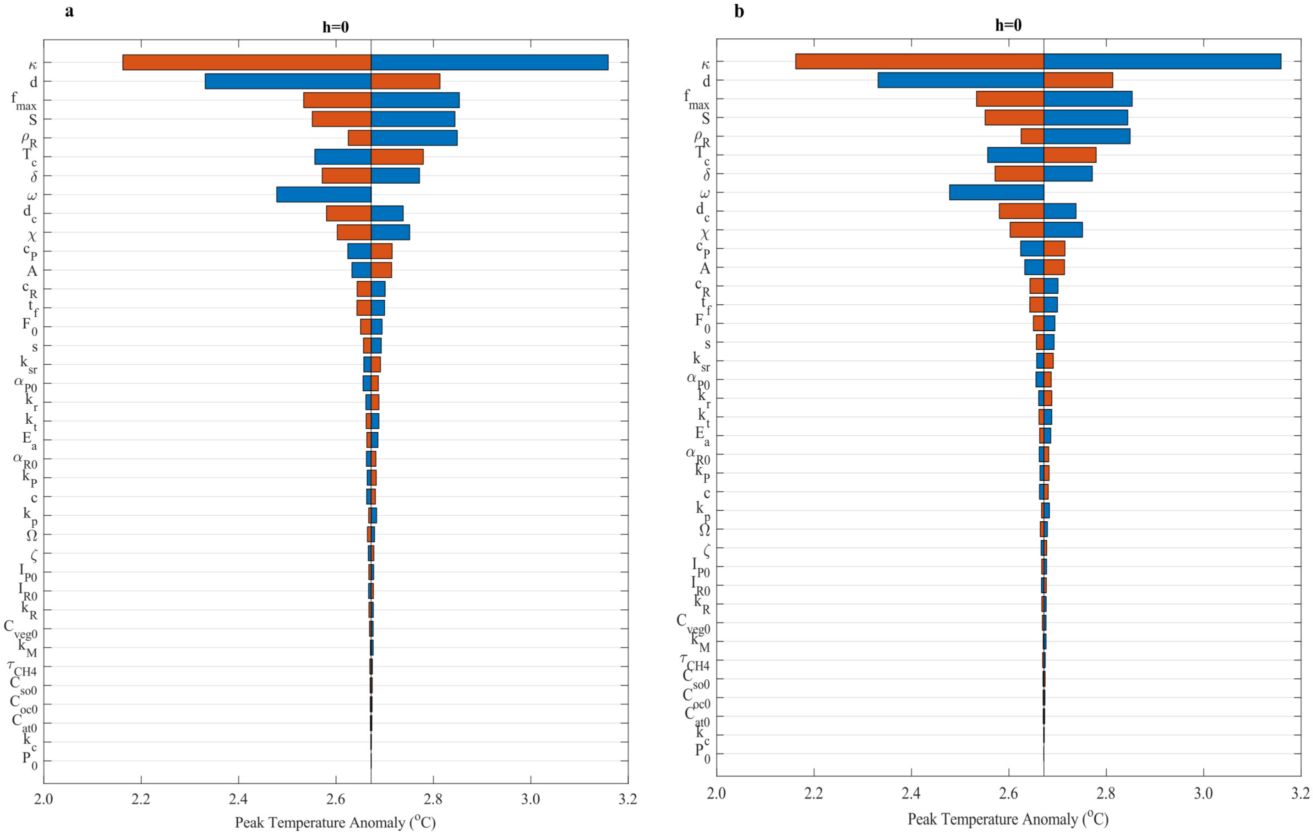


Figure S3: Comparison of (a) no normalization, and (b) social norms normalized, at zero homophily.

There does not seem to be any difference between the main text sensitivity analysis at $h = 0$ and the normalized social norms version sensitivity analysis at $h = 0$.

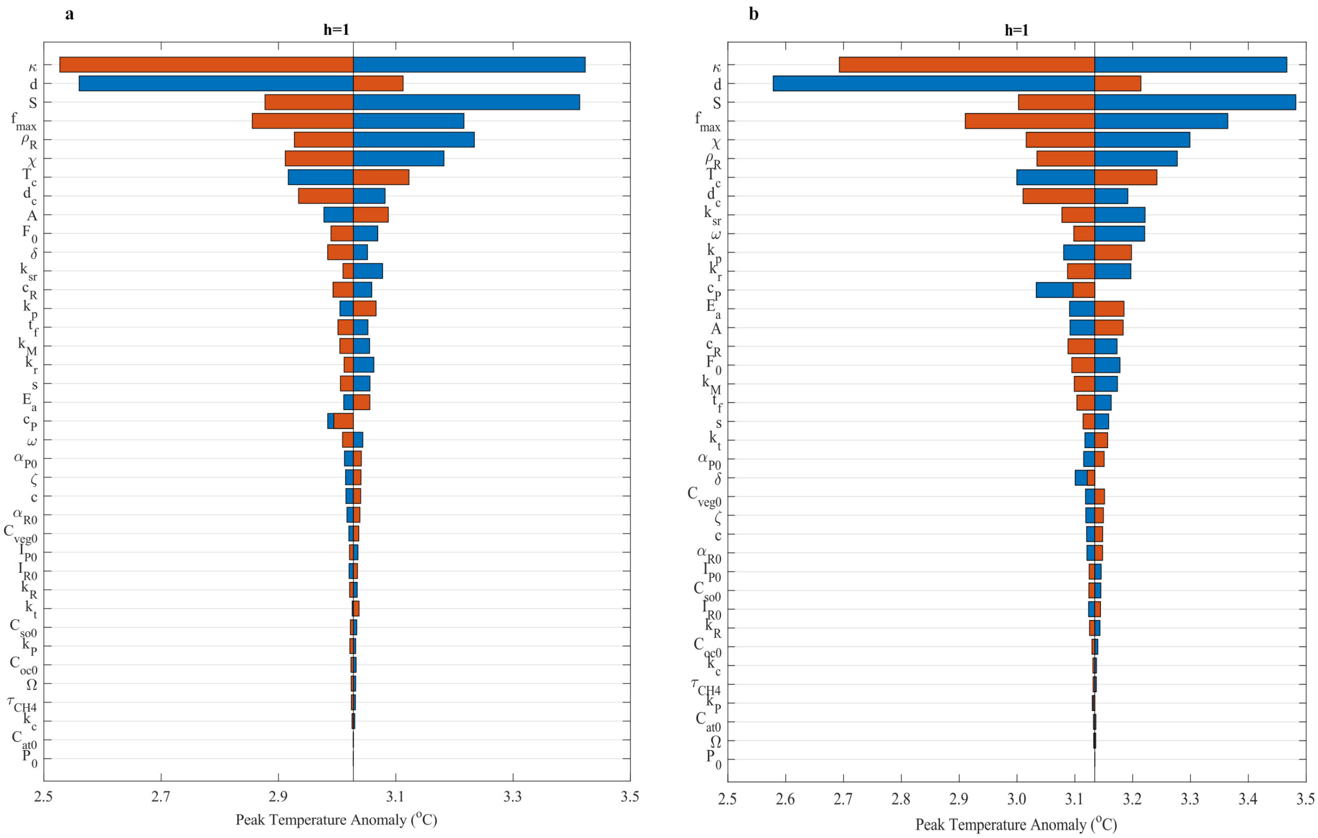


Figure S4: Comparison of (a) no normalization, and (b) social norms normalized, at full homophily.

Compared to the main text's $h = 1$ tornado plot, $\delta$ drops in relative impact, and the peak temperature anomaly at baseline increases by $\approx 1^oC$

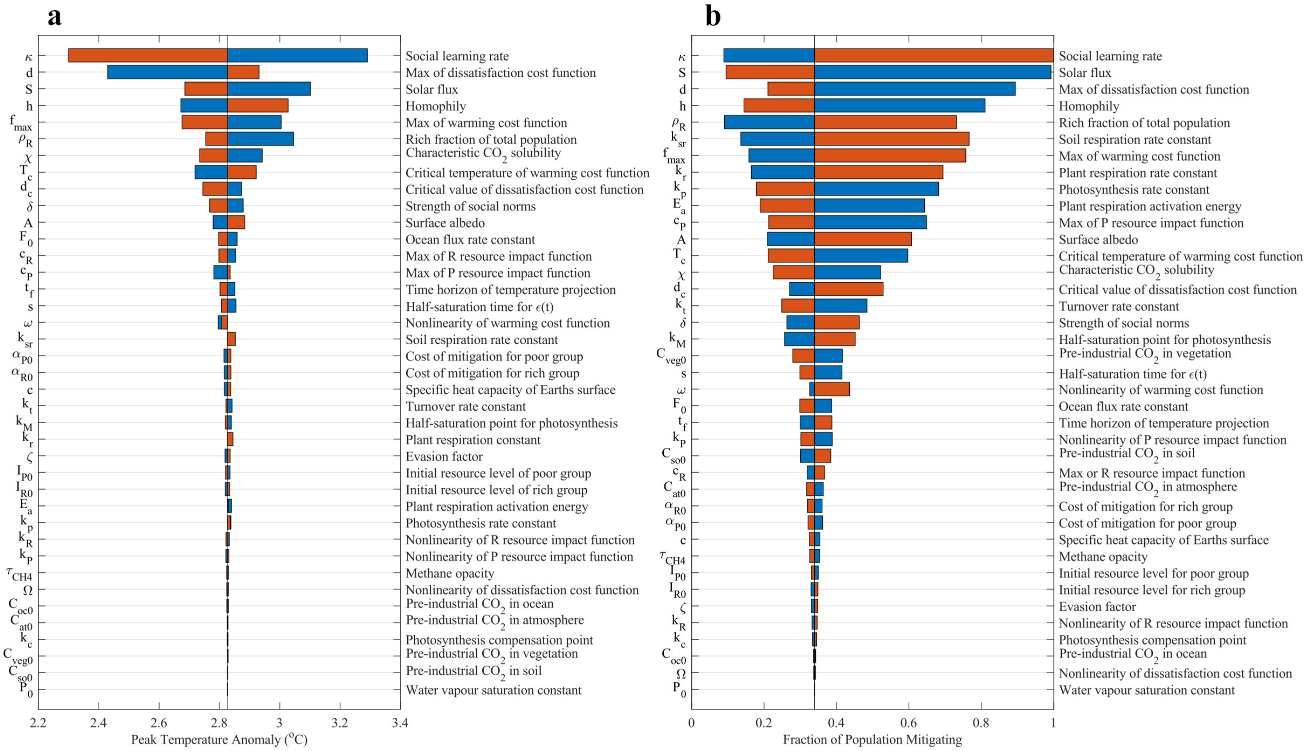


Figure S5: **Climate and social outcomes are most sensitive to uncertainties in social dynamics**. Tornado plots showing the effect on (a) peak temperature anomaly, and (b) fraction of the total population mitigating, when parameters are varied individually. Red (blue) colour denotes the parameter takes its upper (lower) value. Parameters are then ordered by relative impact on peak temperature anomaly. Specific values for each parameter are given in the supplementary information (SI Text Table 2). (b) depicts the mitigation population at year 2100.

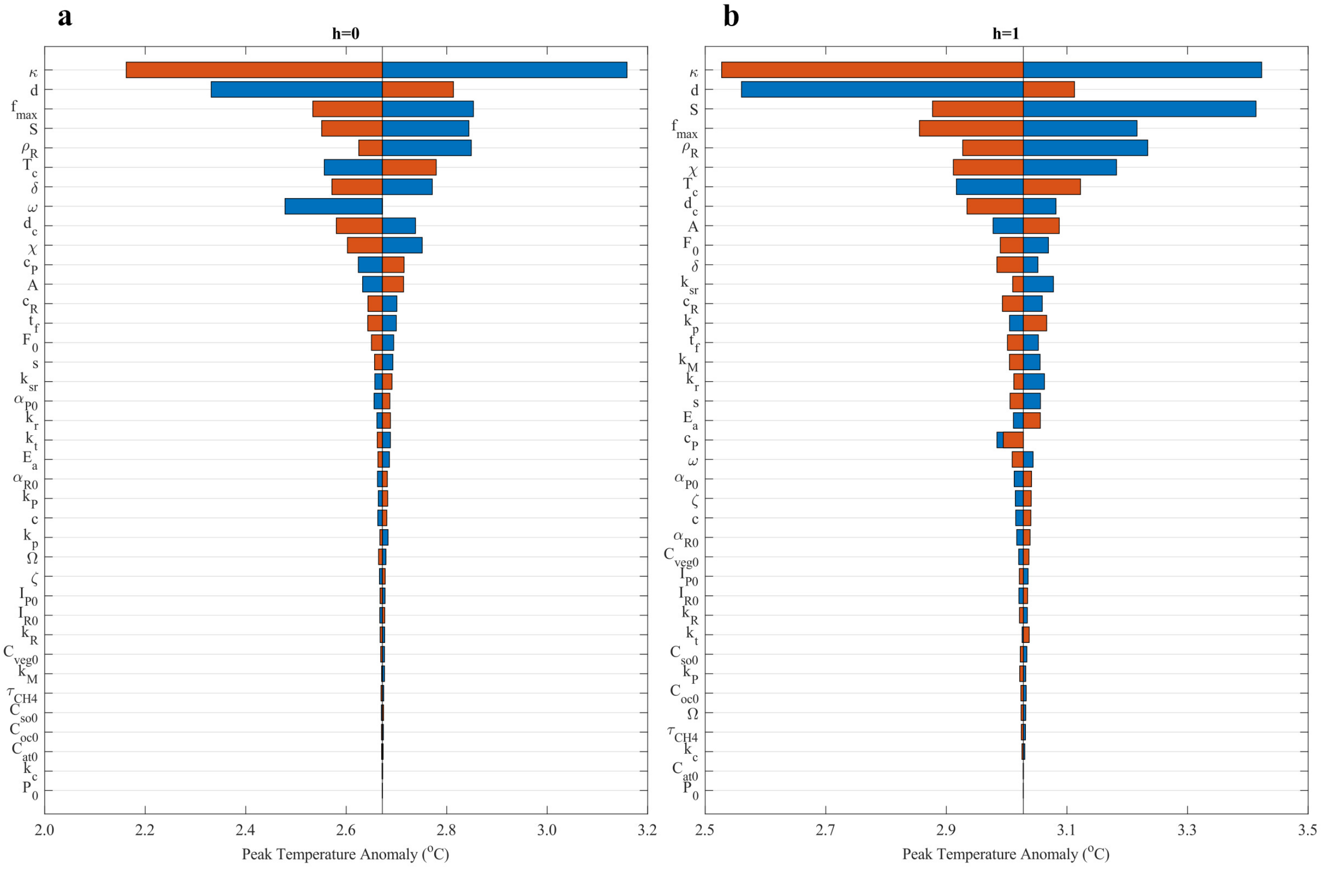


Figure S6: **Homophily extremes change parameter sensitivities**. Tornado plot for (a) no homophily and (b) complete homophily. Red (blue) bar indicates the parameter is taking its upper (lower) value, while all other parameters are at their respective baselines. Parameters are ordered by relative impact from top to bottom. Specific values for each parameter are given in Table 2 (SI Text).

### 1.6 Comparing $\rho_R$ and homophily with dissatisfaction turned off and on

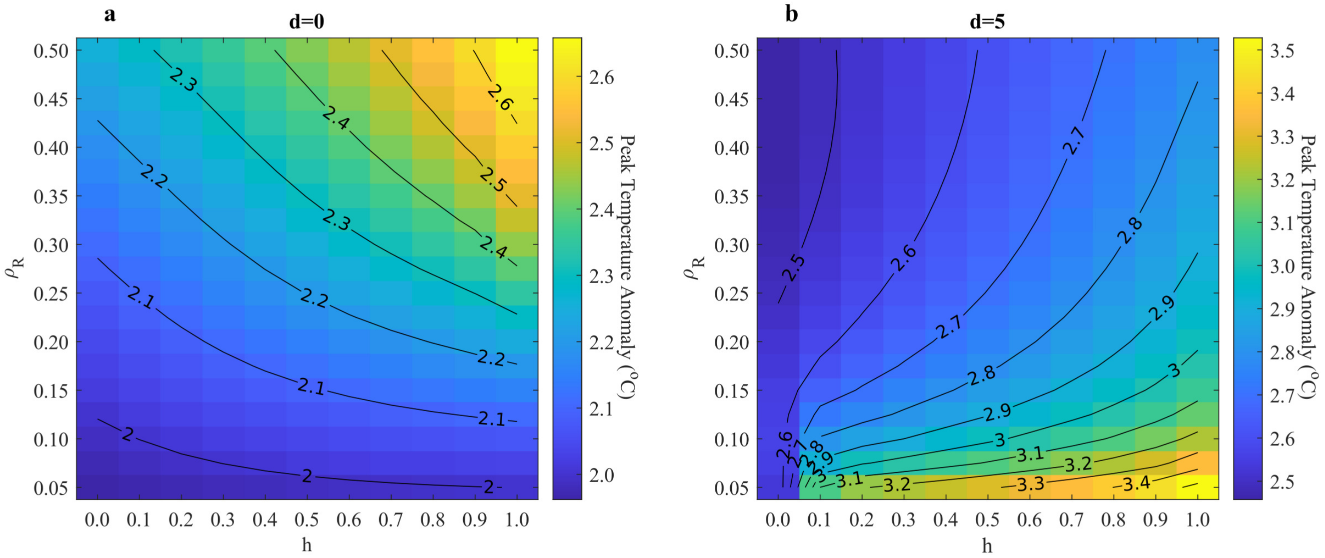


Figure S7: Contour plots showing peak temperature anomaly attained at specific values of homophily ($h$) and proportion of total population in the rich group ($\rho_R$) when (a) dissatisfaction is turned off ($d = 0$) and (b) dissatisfaction is at baseline.

### 1.7 Competition between cost of dissatisfaction and cost of climate change

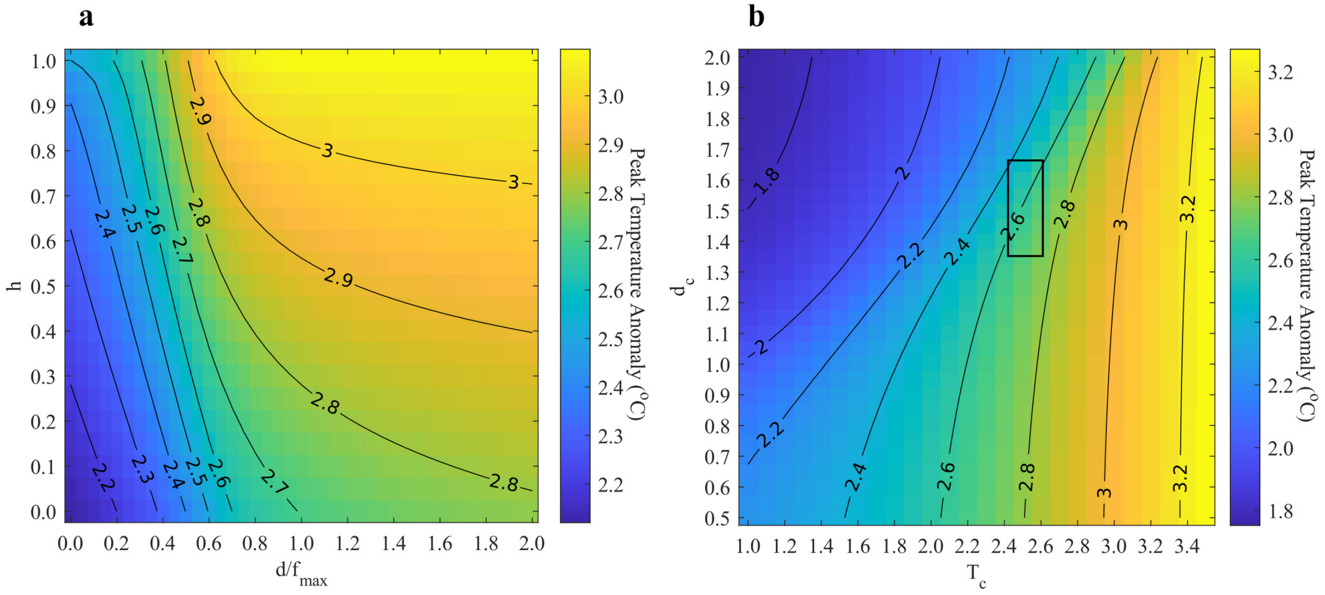


Figure S8: Contour plots showing peak temperature anomaly attained at specific values of (a) homophily ($h$) and relative maximum cost of dissatisfaction to cost of climate change, and (b) critical temperature of $f(T_f)$ vs. critical value of dissatisfaction cost. All other parameters are held at baseline values, as defined in SI Table 1. The window in (b) corresponds to the interval outlined in SI Table 2.

### 1.8 $x_P$-$x_R$ Analysis.

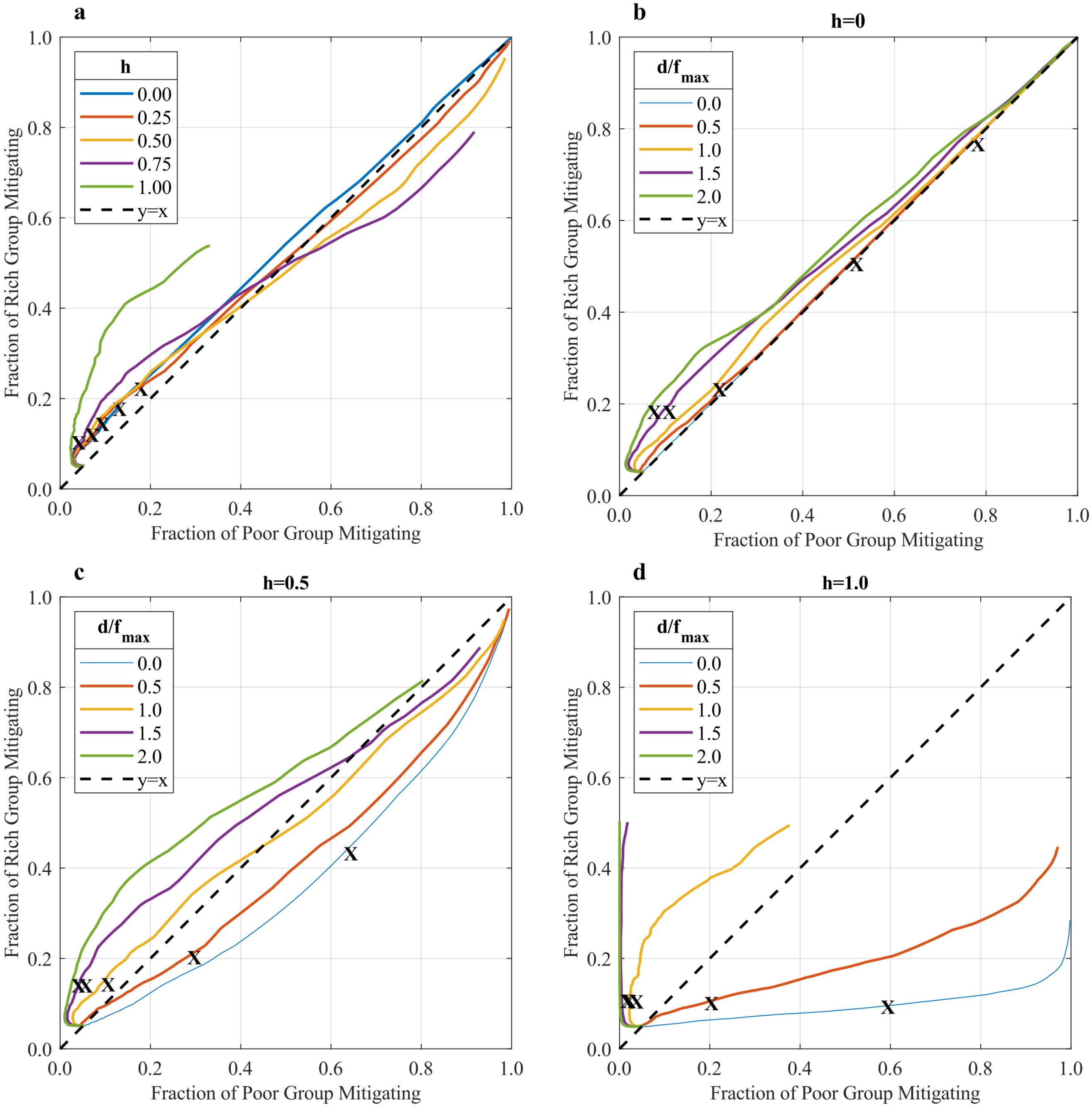


Figure S9: Median proportion of each group mitigating after 100 simulations under (a) changing levels of homophily; (b) zero homophily, and changing levels of maximum cost of dissatisfaction ($d$) relative to the maximum cost of the warming function ($f_{max}$); (c) half homophily, and changing levels of $\frac{d}{f_{max}}$; (d) full homophily, and changing levels of $\frac{d}{f_{max}}$. All other parameters being sampled from triangular distributions defined in Table 1. "X"s mark $t = 2050$, while lines end at $t = 2100$. Note how increasing homophily, and increasing strength of dissatisfaction, both slow the adoption of mitigative behaviour.

#### 1.9 Dependence of peak temperature anomaly plateau on year at which rich group becomes mitigative.

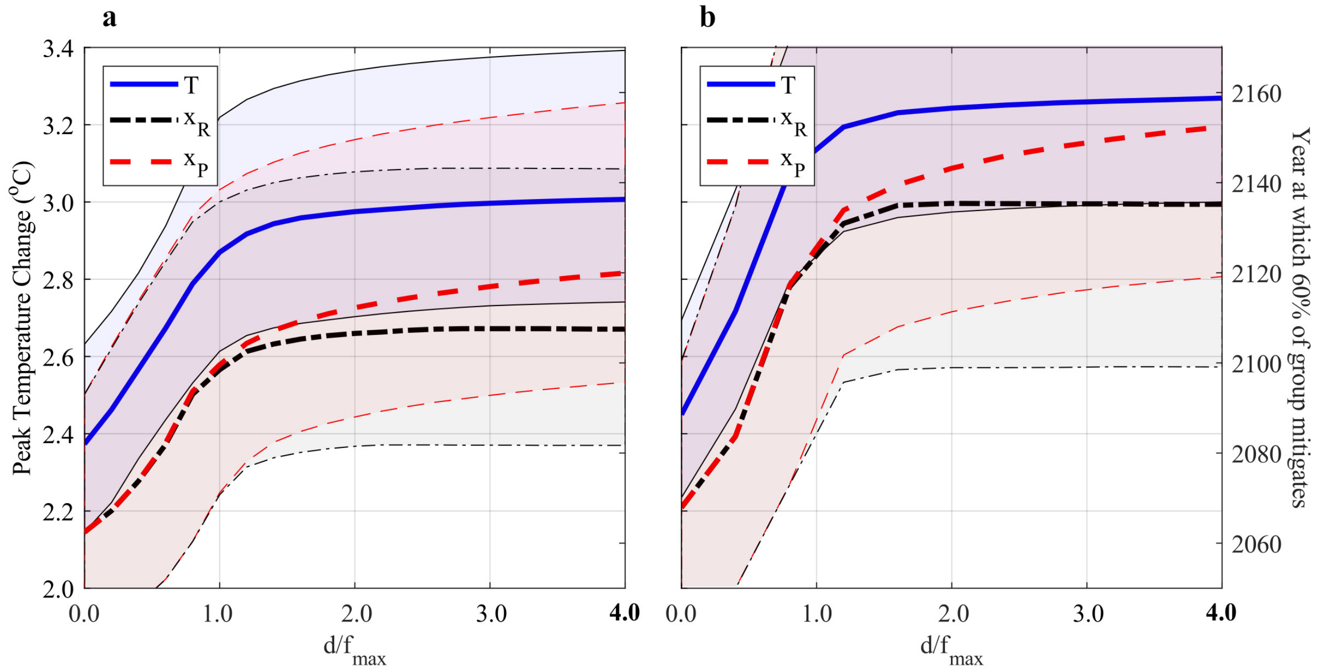


Figure S10: **Peak temperature anomaly plateaus with increasing $\frac{d}{f_{max}}$ because the ratio no longer affects when the rich group becomes mitigative**. Peak temperature anomaly (left y-axis) and year at which 60% of each group is mitigating (right y-axis) vs. relative importance of dissatisfaction and climate change impacts. Median and 95% CIs are calculated over 100 simulations with parameters drawn from triangular distributions with lower and upper bounds defined as: (a) SI Text Table 1; and (b), SI Text Table 2 except $\alpha_{R0}$ is shifted to baseline 1.5. Both panels have fixed $f_{max} = 5$ and $h = 0$.

When $\frac{d}{f_{max}}$ continues increasing above 1, a plateau in peak temperature anomaly, and in year at which 60% of the rich group becomes mitigative, appears. The latter is because $d$ does not directly impact the utility functions of the rich group. The former is because the rich group plateaus, an increase in $d$ does not further delay the rich group from becoming mitigative, leading them to reduce their carbon emissions at the same rate at each value of $\frac{d}{f_{max}}$ above 1. In addition, raising the cost of mitigation for the rich group – increasing $\alpha_{R0}$ from 0.5 (panel a) to 1.5 (panel b) – raises both the peak temperature anomaly and the year at which 60% of the rich group is mitigative; the peak temperature increases by $\approx 0.25^oC$ while the year 60% of the rich group becomes mitigative increases by $\approx 30yrs$

#### 1.10 Model Components

The full social-climate model is given here:

$$\frac{\mathrm{d}}{\mathrm{d}t}x_R(t) = \kappa x_R(t) y_R(t) \left[E_R(T;M) - E_R(T;N)\right] + (1-h)\kappa\left\{ \left[x_P(t) y_R(t) \max\left(E_P(T;M) - E_R(T;N),\ 0\right)\right] - \left[x_R(t) y_P(t) \max\left(E_P(T;N) - E_R(T;M),\ 0\right)\right] \right\}$$

$$\frac{\mathrm{d}}{\mathrm{d}t}x_P(t) = \kappa x_P(t) y_P(t) \left[E_P(T;M) - E_P(T;N)\right] + (1-h)\kappa\left\{ \left[x_R(t) y_P(t) \max\left(E_R(T;M) - E_P(T;N),\ 0\right)\right] - \left[x_P(t) y_R(t) \max\left(E_R(T;N) - E_P(T;M),\ 0\right)\right] \right\}$$

$$\frac{\mathrm{d}C_{atm}}{\mathrm{d}t} = \varepsilon(t)\left[\frac{y_R I_R + y_P I_P}{y_{R0} I_{R0} + y_{P0} I_{P0}}\right] - P + R_P + R_S - F_{aw} - F_{ac}$$
$$\frac{\mathrm{d}C_{oc}}{\mathrm{d}t} = F_{oc}$$
$$\frac{\mathrm{d}C_{veg}}{\mathrm{d}t} = P - R_{veg} - L$$
$$\frac{\mathrm{d}C_{so}}{\mathrm{d}t} = L - R_{so}$$
$$c\frac{\mathrm{d}T}{\mathrm{d}t} = \left(F_d - \sigma T^4\right) a_E$$

Climate variables are expressed as deviations from pre-industrial levels. Definitions and intervals are given in SI Text table 2

### Photosynthesis

Carbon uptake from the atmosphere occurs in the following form:

$$P(C_{at}, T) = k_p C_{veg0} k_{MM} \left(\frac{pCO_{2a} - k_c}{K_M + pCO_{2a} - k_c}\right)\left(\frac{(15+T)^2 (25-T)}{5625}\right)$$

for $pCO_{2a} \geq k_c$ and $-15 \leq T \leq 25$, and zero otherwise.
$pCO_{2a}$ is the mixing ratio of $CO_2$ in the atmosphere and is defined as the ratio of moles of $CO_2$ in the atmosphere to the total number of molecules in the atmosphere $k_a$:

$$pCO_{2a} = \frac{f_{gtm}\left(C_{atm} + C_{atm0}\right)}{k_a}$$

where $f_{gtm} = 8.3259 \times 10^{13}$ converts from $gTC$ to moles of carbon; $C_{atm0}$ is the initial level of carbon in the atmosphere. Photosynthesis satisfies Michaelis-Menton kinetics in $pCO_{2a}$. Optimal photosynthesis occurs at $T = 2$ and rates of photosynthesis declines as $T$ increases above 2.

### Respiration

Plant respiration takes the form of:

$$R_{veg}(T, C_{veg}) = k_r C_{veg} k_A e^{-\frac{E_a}{R(T+T_0)}}$$

This increases with the amount of carbon present in the vegetation, and also with temperature. A positive feedback loop then results with increasing carbon levels. Soil respiration is analogous:

$$R_{so}(T, C_{so}) = k_{sr} C_{so} k_B e^{-\frac{308.56}{T+T_0+227.13}}$$

#### Turnover

In this climate model we assume a constant fraction of plants die over a given unit of time:

$$L(C_{veg}) = k_t C_{veg}$$

The carbon from the dead plants is then fed into the soil reservoir.

#### Ocean Flux

$CO_2$ from the atmosphere transfers to the ocean in the following way:

$$F_{oc}(C_{at}, C_{oc}) = F_0 \chi \left( C_{at} - \zeta \frac{C_{at0}}{C_{oc0}} C_{oc} \right)$$

$\chi$ is the characteristic solubility of $CO_2$ in water and $\zeta$ is the evasion factor [42]. More complex ocean-atmosphere models couple these parameters to chemical dynamics occurring within the ocean itself [29]; however, we find good agreement when comparing this simplification to the full model of [29].

#### Atmospheric Dynamics

Atmospheric dynamics are modelled using the grey-atmosphere approximation presented in [29]. This method models changes in global average temperature using changes in albedo ($A$), incoming solar flux ($S$), and the opacity of $CO_2$, $H_2O_{(}v)$, and $CH_4$. The net downward flux of solar radiation absorbed at the planet's surface is given by:

$$F_d = \frac{(1-A)S}{4} \left( 1 + \frac{3}{4}\tau \right)$$

$\tau$ is the sum of opacities of $CO_2$, $H_2O_{(}v)$, and $CH_4$; each gas' opacity is given by:

$$\tau(CO_2) = 1.73(pCO_2)^{0.263}$$
$$\tau(H_2O) = 0.0126 \left( H P_0 e^{-\frac{L}{RT}} \right)^{0.503}$$
$$\tau(CH_4) = 0.0231$$

with $pCO_2$ being the mixing ratio of $CO_2$ in the atmosphere (defined above), $H$ is the relative humidity, $P_0$ is the water vapor saturation constant, $L$ is the latent heat per mole of water, and $R$ is the molar gas constant.

### 1.11 Comparison of ESM with CMIP5

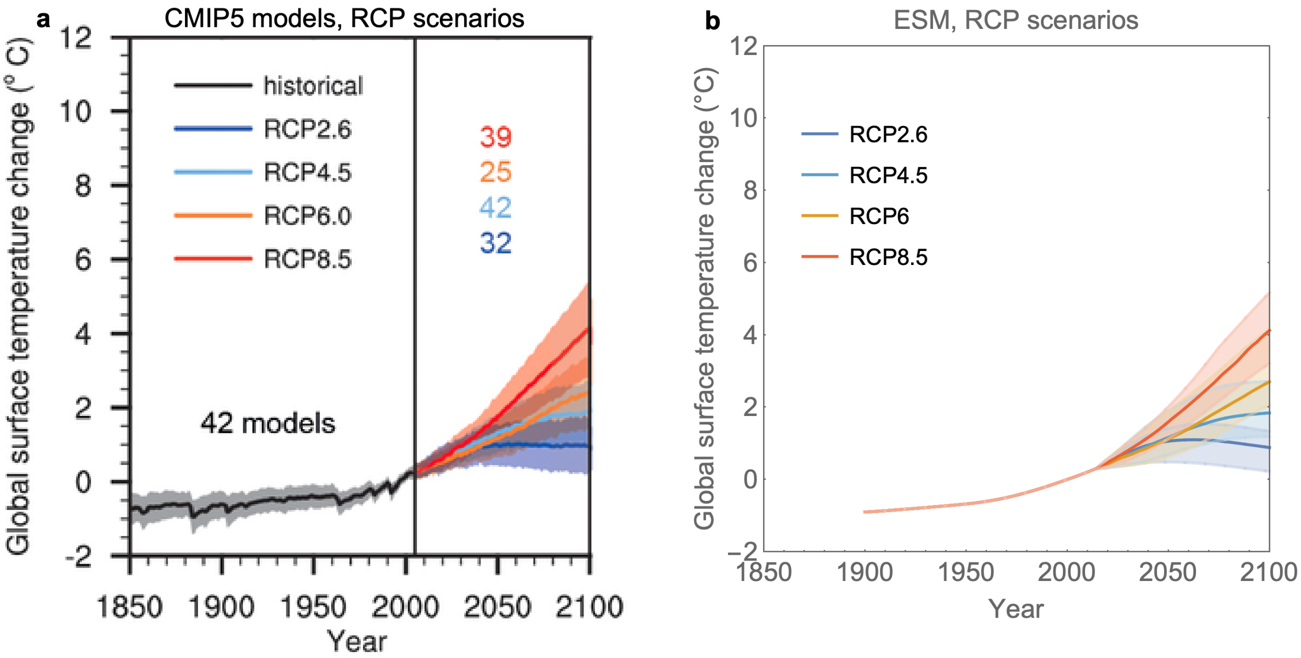


Figure S11: **Comparison of CMIP5 models and ESM:** here we are comparing a simple Earth System model that captures the carbon cycle plus some biochemical processes [29], we can see that it results in very similar temperature trajectories. Obtained from [28]: **a**. is obtained from the IPCC Fifth Assessment Report [12]; **b**. is obtained from [28] and is the result of an ensemble of simulations from the Earth System Model used here and in [28] driven with the same emissions scenarios as the RCPs. Parameters were drawn from triangular distributions with upper and lower bounds given in section 2

### 1.12 Derivation of imitation dynamics

In order to investigate group dynamics coupled to a climate model, we need to know how the prevalence of each strategy between groups evolves through time where strategy change occurs due to an imitation process. We would like to know the imitation dynamics in 2-group asymmetric games; i.e. our goal is to have an imitation dynamic for intra- and intergroup imitation processes. Starting from Eq. 10.10 in [35]:

$$\frac{\mathrm{d}}{\mathrm{d}t} P_a(i,t) = \sum_{i'} \left[ w^a(i|i';t) P_a(i',t) - w^a(i'|i;t) P_a(i,t) \right] \tag{9a}$$

$$w^a(i|i';t) := w_a(i|i';t) + \nu^a(i|i';t) R^a(i|i';t) \tag{9b}$$

$$\nu^a(i,i',t) := \sum_b \nu_{ab}(t) \left[ (f^1_{ab} - f^2_{ab}) P_b(i,t) + (f^2_{ab} - f^3_{ab}) P_b(i',t) + f^3_{ab} \right] \tag{9c}$$

Where Eq. 9a is the master equation for strategy $i$ over time in group $a$. Eq. 9b are the transition rates, with the first term being a spontaneous transition rate (which we will quickly be setting to zero) and the second term corresponding to transitions upon encountering another individual. Lastly, Eq. 9c are rates for the frequency of interactions of an individual from group $a$ with other individuals. Here $f^k_{ab}$ denotes an interaction process of type $k$ between pair $a$, $b$. There are 3 types: (1) imitative, (2) avoidance, and

(3) consensus. We are interested in $k = 1$, an imitative process; so following Helbing in [35] we set:

$$f_{ab}^2 = 0 = f_{ab}^3 \tag{10}$$

However, unlike [35], we do not assume that individuals from distinct groups will not imitate each other; i.e. we do not assume $f_{ab}^1 \approx 0$ for $a \neq b$. Instead we take $f_{ab}^1 = 1$. Following (10) and (9c) we get:

$$\nu^a(i, i', t) := \sum_b \nu_{ab}(t) P_b(i, t) \tag{11}$$

we follow [28, 35] and define our readiness to give up strategy $i$ in favour of strategy $j$ as:

$$R^a(j|i;t) = \max(E_a(j, t) - E_a(i, t), 0) \tag{12}$$

however, we are going to modify it slightly to permit imitation of a strategy from an individual of another group and introduce a homophily (or polarization) parameter $h \in [0, 1]$, controlling the extent to which distinct groups will listen to each other (inspired by [39]). Hence we have

$$R^a(j|i;t) = (1 - h(1 - \delta_{ab})) \max\big(E_b(j, t) - E_a(i, t), 0\big) \tag{13}$$

Where the Kronecker delta ensures that the homophily parameter does not affect interactions between pairs from the same group. Inserting (11) and (13) into (9b) leads to:

$$w^a(i|i';t) := w_a(i|i';t) + \sum_b \nu_{ab}(t) P_b(i, t)\,(1 - h(1 - \delta_{ab})) \max\big(E_b(i, t) - E_a(i', t),\ 0\big) \tag{14}$$

since the first term $w_a(i|i';t)$ denotes spontaneous transitions from strategy $i' \to i$, we will set it to zero because we are focusing only on imitative processes. Our transition rates are then

$$w^a(i|i';t) := \sum_b \nu_{ab}(t) P_b(i, t)\,(1 - h(1 - \delta_{ab})) \max\big(E_b(i, t) - E_a(i', t),\ 0\big) \tag{15}$$

where $b$ runs over all groups including $a$. We can insert (15) into (9a) and get:

$$\frac{\mathrm{d}}{\mathrm{d}t} P_a(i, t) = \sum_{i'} \Bigg\{ \Bigg[ \sum_b \nu_{ab}(t) P_b(i, t)\,(1 - h(1 - \delta_{ab})) \max\big(E_b(i, t) - E_a(i', t),\ 0\big) \Bigg] P_a(i', t) - \Bigg[ \sum_b \nu_{ab}(t) P_b(i', t)\,(1 - h(1 - \delta_{ab})) \max\big(E_b(i', t) - E_a(i, t),\ 0\big) \Bigg] P_a(i, t) \Bigg\}$$

Since the $b$-index is the same for both summations over $b$ we can write:

$$\frac{\mathrm{d}}{\mathrm{d}t}P_a(i,t) = \sum_{i'}\sum_{b}(1-h(1-\delta_{ab}))\Bigg\{\Big[\nu_{ab}(t)P_b(i,t)\max\big(E_b(i,t)-E_a(i',t),\ 0\big)\Big]\,P_a(i',t) - \tag{16}$$

$$\Big[\nu_{ab}(t)P_b(i',t)\max\big(E_b(i',t)-E_a(i,t),\ 0\big)\Big]\,P_a(i,t)\Bigg\} \tag{17}$$

If we split (16) into two terms, one when $b = a$ and one summing over all $b \neq a$, we get

$$\begin{aligned}\frac{\mathrm{d}}{\mathrm{d}t}P_a(i,t) =& \sum_{i'}\Bigg\{\Big[\nu_{aa}(t)P_a(i,t)\max\big(E_a(i,t)-E_a(i',t),\ 0\big)\Big]\,P_a(i',t) - \\ &\Big[\nu_{aa}(t)P_a(i',t)\max\big(E_a(i',t)-E_a(i,t),\ 0\big)\Big]\,P_a(i,t)\Bigg\} + \\ &(1-h)\sum_{i'}\sum_{b(\neq a)}\Bigg\{\Big[\nu_{ab}(t)P_b(i,t)\max\big(E_b(i,t)-E_a(i',t),\ 0\big)\Big]\,P_a(i',t) - \\ &\Big[\nu_{ab}(t)P_b(i',t)\max\big(E_b(i',t)-E_a(i,t),\ 0\big)\Big]\,P_a(i,t)\Bigg\}\end{aligned}$$

simplifying we can get:

$$\begin{aligned}\frac{\mathrm{d}}{\mathrm{d}t}P_a(i,t) =& \sum_{i'}\nu_{aa}(t)P_a(i,t)P_a(i',t)\,\big[E_a(i,t)-E_a(i',t)\big]\ + \\ &(1-h)\sum_{i'}\sum_{b(\neq a)}\nu_{ab}(t)\Bigg\{\Big[P_b(i,t)\max\big(E_b(i,t)-E_a(i',t),\ 0\big)\Big]\,P_a(i',t)\ -\Big[P_b(i',t)\max\big(E_b(i',t)-E_a(i,t),\ 0\big)\Big]\end{aligned}$$ [illegible] (18)

The first term simplifies because $\max(A-B,0)-\max(B-A,0) = A-B$. In the case where we set $h = 1$ (full homophily) then we retrieve the generalized game dynamical equations of [35] (without spontaneous transitions) and have:

$$\begin{aligned}\frac{\mathrm{d}}{\mathrm{d}t}P_a(i,t) &= \nu_{aa}(t)P_a(i,t)\sum_{i'}P_a(i',t)\big(E_a(i,t)-E_a(i',t)\big) \\ &= \nu_{aa}(t)P_a(i,t)\left(E_a(i,t)-\sum_{i'}E_a(i',t)P_a(i',t)\right)\end{aligned}$$

this is because in [35] the assumption of $h = 1$ is assumed in the readiness to change strategy. If we further restrict to two strategies and one population, then we retrieve the DE used in [28]:

$$\frac{\mathrm{d}}{\mathrm{d}t}P_a(i,t) = \nu_{aa}(t)P_a(i,t)(1-P_a(i,t))\big(E_a(i,t)-E_a(i',t)\big)$$

where this can be made even more explicit by replacing $\nu_{aa}$ by by the social learning parameter $\kappa$ and $P_a(i,t)$ by $x$, leading to

$$\frac{\mathrm{d}x}{\mathrm{d}t} = \kappa x(1-x)\big(E(M) - E(N)\big)$$

If instead, starting from Eq.18 we make the choices that there are two groups, $a$ and $b$ (or rich and poor), two strategies – mitigation (M) and non-mitigation (N) – along with with the groups obtaining different payoffs for the same strategies, we obtain Eq. 2 in the main text. This can be made explicit by making the replacements:

$$\begin{aligned} i &= M \\ i' &= N \\ P_a(i,t) &= x_P(t) \\ P_a(i',t) &= y_P(t) \\ P_b(i,t) &= x_R(t) \\ P_b(i',t) &= y_R(t) \\ \nu_{aa} = \nu_{ab} &= \kappa \end{aligned}$$

giving

$$\begin{aligned} \frac{\mathrm{d}x_P}{\mathrm{d}t} =& \kappa x_P y_P \left[E_P(M) - E_P(N)\right] + \\ (1-h)\kappa & \Big\{ \left[x_R \max\big(E_R(M) - E_P(N),\ 0\big)\right] y_P - \left[y_R \max\big(E_R(N) - E_P(M),\ 0\big)\right] x_P \Big\} \end{aligned} \tag{19}$$

Note that we made the choice $\nu_{aa} = \nu_{ab} = \kappa$, meaning social learning occurs at the same rate between groups as within groups.
